# Brainstem development requires galactosylceramidase and is critical for the pathogenesis of Krabbe Disease

**DOI:** 10.1101/2020.03.25.007542

**Authors:** Nadav I. Weinstock, Conlan Kreher, Jacob Favret, Ernesto Bongarzone, Lawrence Wrabetz, M. Laura Feltri, Daesung Shin

## Abstract

Krabbe disease (KD) is caused by a deficiency of galactosylceramidase (GALC), which induces demyelination and neurodegeneration due to accumulation of cytotoxic psychosine. Hematopoietic stem cell transplantation (HSCT) improves clinical outcomes in KD patients only if delivered pre-symptomatically. We hypothesized that the restricted temporal efficacy of HSCT reflects a requirement for GALC in early brain development. Using a novel *Galc* floxed allele, we induced ubiquitous GALC ablation (*Galc*-iKO) at various postnatal timepoints and identified a critical period of vulnerability to GALC ablation between P4-6. Early *Galc*-iKO induction caused a worse KD phenotype, higher psychosine levels, and a significantly shorter life-span. Intriguingly, GALC expression peaks during this critical developmental period. Further analysis revealed a novel cell autonomous role for GALC in the development and maturation of immature T-box-brain-1 positive brainstem neurons. These data identify a perinatal developmental period, in which neuronal GALC expression influences brainstem development that is critical for KD pathogenesis.

## Introduction

Krabbe disease (KD) is a demyelinating and neurodegenerative lysosomal storage disorder (LSD), often fatal in infancy^1^. KD is caused by mutations in GALC, a galactolipid hydrolase that must be trafficked to the lysosome for proper functioning^2^. GALC is involved in the normal turnover of myelin by hydrolyzing galactosylceramide, a major sphingolipid constituent of myelin important in myelin compaction^3^. Unlike other LSDs, the primary substrate of GALC, galactosylceramide, does not accumulate broadly in KD tissues. Instead, a minor substrate of GALC, psychosine (galactosylsphingosine), accumulates to toxic levels, causing extensive demyelination and the bulk of KD pathological findings (‘psychosine hypothesis’^4^). In support of this hypothesis, *in vitro* studies show that treating cells with psychosine increases proapoptotic factors and kills oligodendrocytes (OL)^5^, Schwann cells^6, 7^ and neurons^8^. The long-held psychosine hypothesis was recently confirmed, *in vivo*, which demonstrated that the abnormal accumulation of psychosine is toxic and is generated catabolically through the deacylation of galactosylceramide by acid ceramidase^9^

More than 85% of KD patients exhibit the rapidly progressive infantile-onset form of disease, which leads to death by two years of age. Although there is no cure for Krabbe disease, hematopoietic stem cell therapy (HSCT) attenuates neurologic deterioration and improves developmental gains^10^. These benefits are particularly sensitive to the severity of disease at transplantation, and are only beneficial if delivered at a clinically defined ‘pre-symptomatic’ timepoint^10^. Intriguingly, pre-clinical gene therapy trials in *twitcher* mice are also time-sensitive, and have been shown to be more efficacious if delivered shortly after birth^11^. These data therefore suggest a pre-symptomatic ‘therapeutic window’, in which treatment of KD with HSCT or gene therapy is more efficacious. Why treatment must occur so early remains unknown. While GALC should intuitively be required during myelination, many of these temporal events seem to precede the bulk of CNS myelination. Furthermore, patients diagnosed *in utero*, and treated within the first few weeks of life, do better than patients treated at 1-2 months old. These findings underlie the necessity for properly defining the precise timepoint in which GALC is first required.

In a previous study, we used a global metabolomic analysis of *twitcher* hindbrains to detect the earliest biochemical changes that occur in KD pathogenesis. While we saw many metabolic changes at an early symptomatic stage of disease (P22), we were surprised to find a number of biochemical processes that were also significantly altered at a ‘pre-symptomatic’ timepoint (P15)^12^. These changes, though subtle, reflected diverse neuro-metabolic functions relating to glycolysis, the pentose phosphate pathway, hypoxanthine metabolism and mannose-6-phosphate, an important residue involved in lysosomal enzyme trafficking^12^. These findings, along with the observed pre-symptomatic ‘therapeutic window’, led us to hypothesize that GALC has important and specific functions in early brain development.

Several ubiquitous *Galc* mutant mice have been used to study KD, including *twitcher*^13^, *twi-5J*^14^, humanized *GALC* transgenic^15^, *GALC-Gly270Asp*^16^, *Galc-His168Cys* knock-in mice^17^ and Saposin A knockout mice^18^. Although these models were instrumental in the characterization of KD and the development of various therapy strategies, none of them could identify the temporal effect of GALC deficiency on the progression of KD. We therefore engineered a conditional *Galc* floxed mouse by gene targeting, which provided us with the opportunity to directly ask at which age is *Galc* required. Both constitutive *Galc*-knockouts (KO) and induced *Galc*-KOs (iKO) generated from the conditional floxed allele recapitulated a range of neurologic features seen in KD patients. Our study revealed a key developmental process which requires GALC at perinatal period. Induced deletion of *Galc* prior to P4 resulted in severe neurodevelopmental defects that were particularly worse in the brainstem. Conversely, deletion of *Galc* after P6 resulted in significantly enhanced survival and reduced pathology. This study demonstrates that temporal GALC expression is likely the key contributor to brainstem development. Augmenting GALC levels at, or prior to, this newly defined perinatal period would likely improve the efficacy of therapeutic interventions for KD.

## Results

### The *Galc* knockout mouse is an authentic model of KD

To understand the temporal requirements of GALC, and its relationship to the progression of KD, we developed a conditional *Galc* floxed mouse (Supplementary Fig. 1a), maintained on a congenic C57BL/6 background. To determine if these mice could be used to accurately model KD, global *Galc*-KO mice derived from the *Galc* floxed allele were directly compared to the well-studied *twitcher* (*Galc*^*W339X*^) mice^13^. *Galc*-KO mice were generated by crossing *Galc* floxed mice to CMV-Cre, in which Cre is expressed ubiquitously^19^. PCR analysis of total brain genomic DNA revealed that the *Galc* gene was efficiently deleted (Fig. 1a). Northern blot analysis of total brain RNA from *twitcher (Galc*^*twi/twi*^), *Galc*-KO (*Galc*^*-/-*^), and WT littermates (*Galc*^*+/+*^) showed that *Galc* transcripts were entirely removed from *Galc*-KO (Fig. 1b). Like *twitcher*, homozygous *Galc*-KO had no GALC activity (Fig. 1c) and developed the same phenotype as *twitcher*^20, 21^, namely severe motor coordination defects, a reduced life-span of approximately 45 days and attenuated growth beginning at P21 (Fig. 1d-g). Both *Galc-*KO *and twitcher* mice had an increase in markers of brain inflammation such as toll-like receptor 2 (TLR2), CD68/CD163, Iba1 (all detect microgliosis) and glial fibrillary acidic protein (GFAP; astrogliosis) (Fig. 1h-i). Taken together, the novel *Galc*-KO model has a KD-like phenotype similar to *twitcher* and is a true GALC null allele.

**Figure 1.**
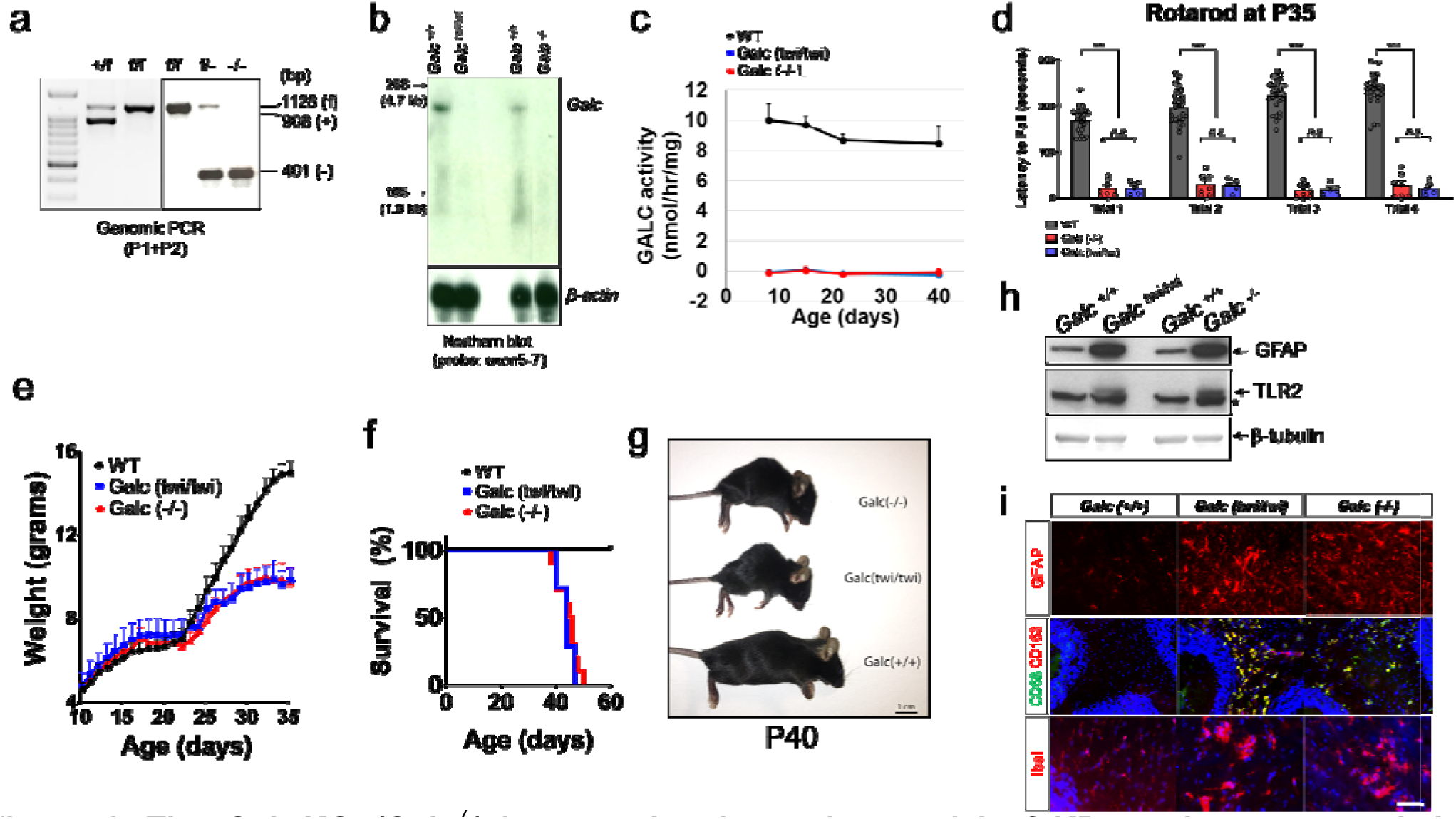
The *Galc*-KO *(Galc*^*-/-*^) is an authentic murine model of KD, analogous to *twitcher* (*Galc*^*twi/twi*^). (**a**) PCR-based genotyping of mice (wild-type [+], conditional [f], and KO [-]) using the P1-P2 primer pair in Supplementary Fig. 1a. *Galc*-KO was generated by mating the *Galc* conditional allele (f) to ubiquitous Cre (CMV-Cre). The floxed locus is efficiently recombined. (**b**) Northern blot analysis of total brain RNAs from P30 *twitcher* (*Galc*^*twi/twi*^), *Galc*-KO (*Galc*^*-/-*^), and wild-type littermate (*Galc*^*+/+*^). The probe spanning exon 5 to 7 of *Galc* cDNA showed that *Galc* transcripts were successfully removed from the *Galc*-KO, similar to the situation in the *twitcher*. (**c**) There were no remaining GALC activities in the brains of both *twitcher* and homozygous *Galc*-KO. (**d**) Rotarod analysis of P35 animals showed that both *twitcher* and *Galc*-KO had the same poor performance. (**e**) Both mice showed a reduced growth curve after P21. (**f**) *Galc*^*-/-*^ survived ∼45 days, like *Galc*^*twi/twi*^. n=10. (**g**) Body size of moribund *Galc*-KO and *twitcher* mice was much smaller than WT. (**h**) Western blot analysis revealed that dramatic increases in the markers of astrocytosis (glial fibrillary acidic protein [GFAP]) and activated microglia (toll-like receptor 2 [TLR2]) in the brain of *Galc*^*-/-*^ as *Galc*^*twi/twi*^ compared to WT *Galc*^*+/+*^. Asterisk (*) is a non-specific band. (**i**) Immunohistochemistry on cryosections of cerebellum white matter showed activated astrogliosis (GFAP) and activated microglia (IbaI, CD68, and CD163), in the brains of both *Galc*^*-/-*^ and *Galc*^*twi/twi*^. Scale bar=100 *µ*m. DAPI is blue-colored. Animals in **g-i** were P40.

### Survival of CAG-Cre/ER^T^ driven *Galc*-iKO mice is dependent on the timing of *Galc* deletion

To determine the age by which *Galc* must be deleted to trigger KD pathogenesis, we crossed *Galc* floxed mice with inducible ubiquitous Cre mice (CAG-Cre/ER^T^)^22^ to generate *Galc* inducible knockout mice *(Galc-*iKO) (Fig. 2a). To maximize the efficiency of *Galc* ablation, we used haplodeficient *Galc*^*flox/null(-)*^ mice. *Galc*-iKO mice were induced between P0 and P10. As expected, perinatal induction of *Galc* ablation (occurring on the day of birth, postnatal day 0) produced a robust neurologic phenotype that closely mirrored the *Galc*-KO. These mice developed a clinical phenotype by P35, including irritability, tremor, wasting, hindlimb paralysis, and ultimately death by P60 (Fig. 2b-c). *Galc*-iKO mice induced between P1-P4 (hereafter ‘*Galc*-iKO≤P4’) exhibited a similar clinical phenotype and also survived to 60 days. Surprisingly, when *Galc* deletion was initiated between P6 and P10 (hereafter ‘*Galc*-iKO≥P6’), the mice developed a significantly protracted clinical course and survived until P90 (Fig. 2b-c). *Galc*-iKO≥P6 had no obvious neurological phenotype or somatic growth defects until around P60 (Fig. 2c; iKO P8). Compared to *Galc*-iKO≤P4, *Galc*-iKO≥P6 had significantly less KD pathology at P60 including fewer inflammatory globoid cells (Fig. 2d-f). These data suggest that GALC ablation at an early developmental timepoint dramatically influences the clinical course of KD and overall degree of pathology produced.

**Figure 2.**
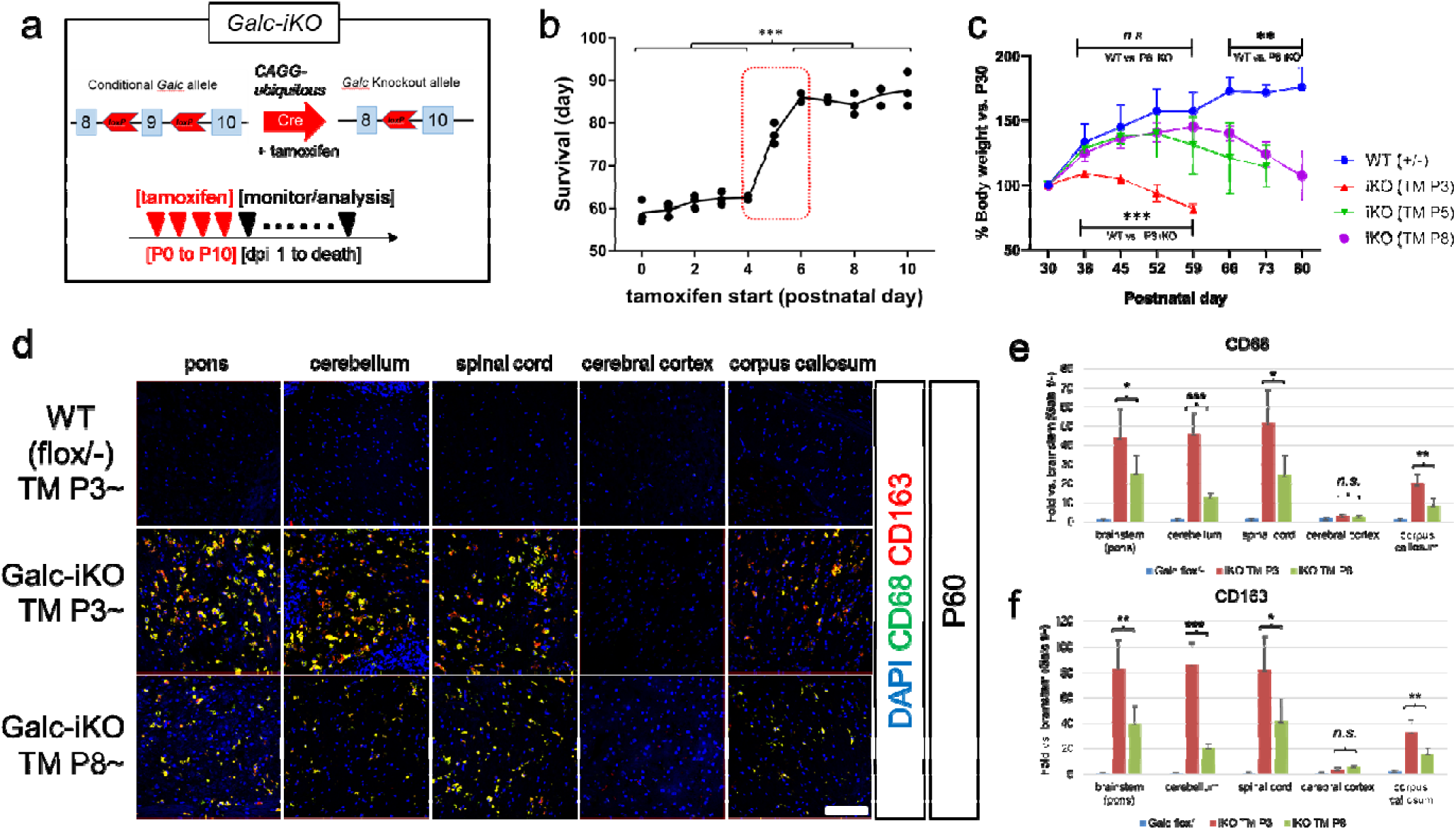
Differential survival of CAG-Cre/ER^T^ driven *Galc*-iKO mice depending on the starting time of tamoxifen injection. (**a**) *Galc* floxed mice were crossed with inducible ubiquitous Cre mice (CAG-Cre/ER^T^), and induced with tamoxifen for four consecutive days. (**b**) When the deletion started at or before P4, the mice only survived about 60 days, but when *Galc* deletion was induced at or after P6, the mice had no phenotype until P60, started to show a tremor after P60 and survived almost 3 months. (**c**)Body weight of *Galc*-induced iKO mice were drastically reduced at before 20-25 days from the last day. n=3. (**d**) The late induced mice (TM starting at P8) were not severe as the early induced *Galc*-iKO (TM at P3), when the latter were moribund at P60. Inflammation markers CD68/CD163 levels were significantly lower in the late induced brains compared to those of the early induced iKO (**e-f**). Scale bar=100 *µ*m.

### The CAG-Cre/ER^T^ transgene recombines efficiently with tamoxifen, regardless of induction time

We were concerned that the differential survival of *Galc*-iKO mice could be due to technical limitations of our inducible system caused by temporal variability in recombination efficiencies. To investigate the temporal recombination efficiency of our system quantitativly, we used a tdTomato Cre reporter^23^. We began by crossing the tdTomato transgene with the CAG-Cre/ER^T^ allele and induced recombination with tamoxifen at P3 (*Galc*-iKO≤P4) or P8 (*Galc*-iKO≥P6). Both timepoints showed an equal number of tdTomato-positive cells and tdTomato protein levels in brain (Fig. 3a-c), suggesting that similar recombination efficiencies. Furthermore, GALC enzymatic activity were similarly absent from the brains of *Galc-*iKO mice induced early and late (Fig. 3d), with only 2% of WT GALC activity remaining. This finding was likewise confirmed by immunofluorescence for GALC expression before and after induction of early and late timepoints (Fig. 3e, f). We also considered that GALC protein could be particularly stable, and could theoretically exist well beyond the induction of *Galc* DNA ablation. We therefore determined the half-life of GALC protein, *in vitro*, and found it to be approximately 3 hours (Fig. 3g), fitting well with the near total reduction in GALC activity seen at both timepoints, *in vivo* (Fig. 3d). These data collectively suggest that technical issues regarding differences in inducing recombination do not likely explain the altered survival and clinical course seen in differential ablation of GALC.

**Figure 3.**
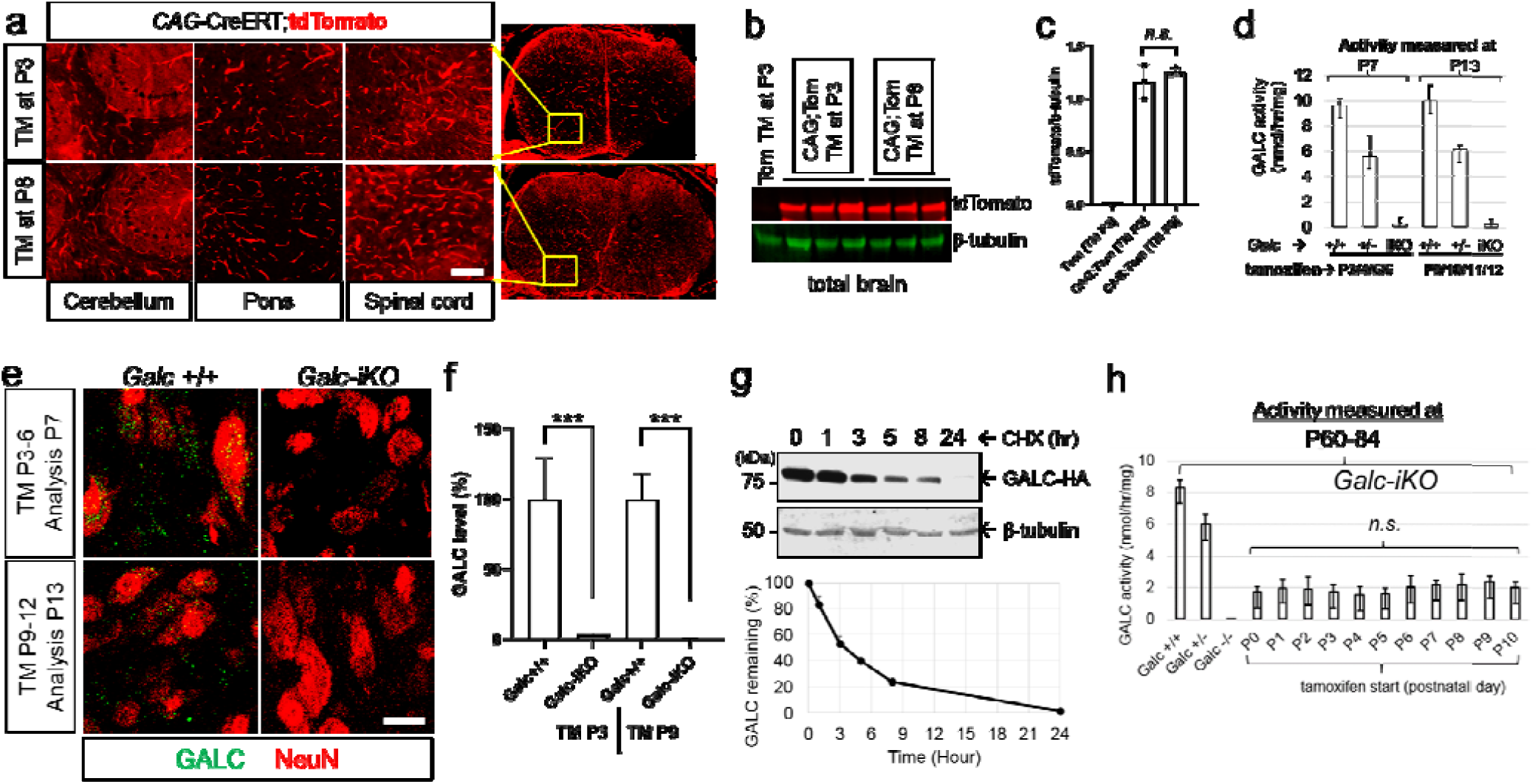
The CAG-Cre/ER^T^ transgene induces recombination efficiently, regardless of induction time. (**a-c**) CAG-Cre/ER^T^;tdTomato mice were induced with tamoxifen starting at P3 or P8, and dissected at P16. The two induction times have a similar number of tdTomato-positive cells in brain and spinal cord (**a**) and the same level of tdTomato protein in whole brains (**b-c**), suggesting no difference in recombination efficiency. Scale bar=100 *µ*m. (**d**) Regardless of induction time, brain GALC activity was efficiently removed 24 hours after tamoxifen injection (from the last day of 4 consecutive days), and were only 2% of normal levels in the total brains. (**e-f**) GALC protein was efficiently depleted 24 hours after tamoxifen injection in the brains of *Galc-*iKO, regardless of induction timing. The image is from pons. Scale bar=20 *µ*m. (**g**) Western blot analysis of HEK-293T cell lysates transfected with HA-GALC, treated with 100 μg/ml cycloheximide, harvested at the indicated time points showed that the half-life of GALC protein, *in vitro*, is about 3 hours. (**h**) **G**ALC activity was not detected at the last day of survival in the whole brains of *Galc*-KO mice (P40), when mice were moribund and paralyzed. Surprisingly, when *Galc*-iKO brains were sampled at similar moribund conditions (between P60 and P84), substantial brain GALC activity had somehow returned and were about 20% of normal levels, regardless of induction starting time. n≥3.

Finally, we asked if the reduction in GALC activity was equally sustained throughout life when induced early or late. There was no significant difference in the residual brain GALC activities of animals within different induction timings. However, we were surprised to find that GALC activity returned to about 20% of WT levels (*Galc*^*+/+*^) in all late-stage, moribund *Galc*-iKO brains (Fig. 3h). Cre/ER^T^ is only active as long as tamoxifen persists^24^. It is possible that the few remaining WT cells proliferate and contribute to restore some levels of GALC activity, which cannot however rescue KD as the mice are terminally sick. In addition, it was particularly intriguing that the residual GALC was unable to cross-correct affected tissues, as deduced from the severe phenotype. This result may correlate with deteriorative moribund pathology of HSCT-treated *twitcher* and KD patients despite the increased GALC activity^25, 26, 27, 28^, emphasizing the importance of providing GALC at a specific developmental period.

### Induction timing of *Galc*-iKO affects the differential accumulation of psychosine

While mice induced after P6 survived longer than those induced earlier, all *Galc*-iKO mice eventually developed a rapidly progressive neurologic decline. Morphometric analysis of optic nerves from early and late-induced moribund *Galc*-iKO mice exhibited similar degrees of demyelination and axonal degeneration (Fig. 4a-c). This was also true for spinal cord tissues, though a more pronounced difference in axonal pathology occurred between early and late induced animals (Fig. 4d-f). Globoid cells and neuroinflammation also accumulated in both end stage *Galc*-iKO≤P4 and *Galc*-iKO≥P6 brains (Fig. 5a-d), though to a slightly higher degree in early induced mice and especially evident in the hindbrain (pons, cerebellum, and spinal cord). Taken together, moribund *Galc*-iKO mice all develop canonical KD pathology that are qualitatively similar to each other, regardless of induction time.

**Figure 4.**
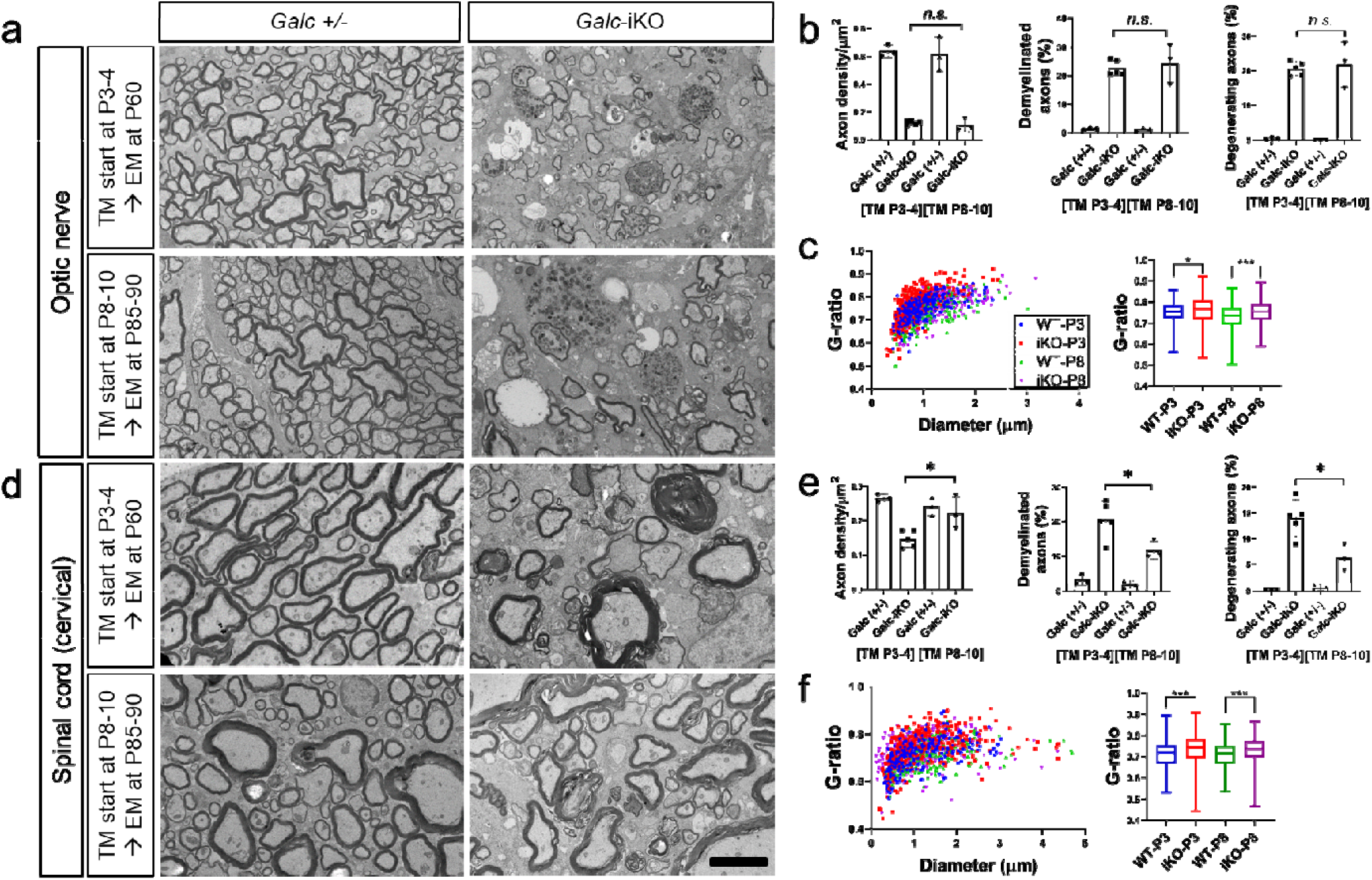
*Galc*-iKO brains have morphological signs of demyelination and axonal degeneration regardless of induction starting time. EM analysis of the optic nerves (**a**) and spinal cords (**d**) and their morphometry-quantifications (**b, e**) from *Galc-*iKO and corresponding WT induced deletion at either starting at P3-4 (*Galc*-iKO≤P4) and P8-10 (*Galc*-iKO≥P6), showed that both conditional *Galc* iKO brains have morphological signs of demyelination, axonal degeneration and gliosis. Although there was no difference of those pathological extents in the optic nerves between the early and late induced *Galc*-iKOs, the morphometric pathological parameters, such as degenerating and demyelinating axons, were slightly less present in the spinal cord of *Galc*-iKO≥P6 than *Galc*-iKO≤P4. Scale bar=4 *µ*m. (**c, f**) The myelin sheaths of both optic nerves and spinal cords were significantly thinner in both *Galc*-iKO≤P4 and *Galc*-iKO≥P6 vs. each WT.

**Figure 5.**
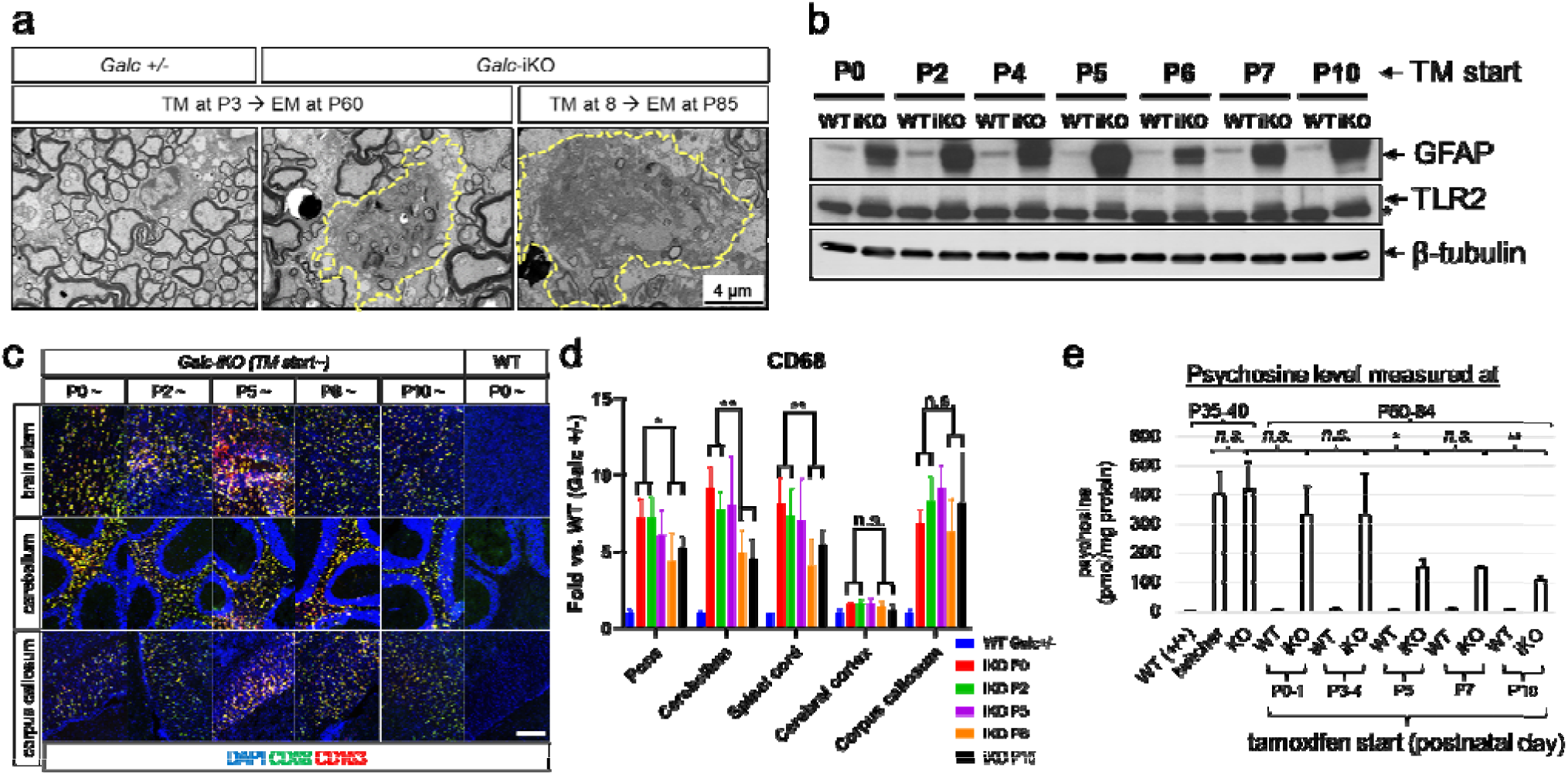
Moribund *Galc*-iKO mice exhibit qualitatively similar KD pathology. Regardless of induction starting time, the moribund *Galc*-iKO mice had the appearance of globoid cells (**a**) and increased of inflammation proteins such as GFAP and TLR2/CD68/CD163 (**b-c**), compared to and WT littermate (*Galc*^*+/+*^). Asterisk (*) is a non-specific band. Scale bar=100 *µ*m. (**d**) Quantitative analysis showed that inflammation marker (CD68) level was slightly lower in the hindbrains of *Galc-*iKO induced after P5 compared to those of the induced before P5. (**e**) Psychosine were measured in the cervical spinal cords of the *Galc-*iKO, sampled when the mice were moribund (P60-84). Although psychosine levels were increased regardless of induction starting time, there was much less psychosine accumulation in the *Galc*-iKO≥P6 compared to the *Galc*-iKO≤P4, indicating of a correlation of psychosine level with the survival.

To elucidate the cause of pathology in moribund mice, we measured psychosine levels in the cervical spinal cord of *Galc*-iKO mice. Psychosine is a highly cytotoxic lipid, capable of inducing cell death of myelinating cells and neurons^8, 29, 30^, and furthermore its accumulation is correlated with severity of disease and age of onset^31^. High performance liquid chromatography tandem mass spectrometry (LC–MS–MS) showed 300-350 pmol of psychosine accumulated per mg of protein in the moribund *Galc*-iKO≤P4. This was far higher than the psychosine concentration in control spinal cords (5-8 pmol/mg of protein, Fig. 5e). Although psychosine also accumulated highly in *Galc*-iKO≥P6 spinal cords (110-150 pmol/mg of protein), these levels were significantly less than the *Galc*-iKO≤P4. The level of psychosine in *Galc*-iKO induced at P5 was in the middle of these extremes (150-160 pmol/mg of protein), suggesting that the concentration of psychosine is directly correlated with pathology and survival of *Galc*-iKO mice.

### GALC expression reaches its highest peak at P5 in the WT brain

We hypothesized that the differential survival of *Galc*-iKO mice induced at pre-symptomatic perinatal ages may reflect a role for GALC in early brain development. We began to address this point by seeking to better understand how *Galc* expression is regulated during postnatal brain development. To do so, we analyzed the mRNA transcripts, protein and activity of GALC in WT brains. Interestingly, *in situ* hybridization and Northern blot analyses showed that *Galc* transcript levels are low at P0, but increase and reach their highest peak at P5 in WT brain (Fig. 6a-b). These peaks correlate closely with the observed clinical worsening and effects on survival that occurred when GALC was ablated temporally (Fig. 2b). Similarly, GALC enzymatic activity mirrored mRNA patterns and peaked at P5 (Fig. 6c), supporting the idea of a developmental process that requires GALC function. Notably, the expression pattern of *Galc* transcripts by *in situ* hybridization suggest *Galc* may also be expressed in non-myelin regions of the P5 brain, particularly in neurons (Fig. 6a and 3e). For example, *Galc* transcripts were clearly expressed in P5 hippocampal neurons and granular neurons of the cerebellum (Fig. 6a enlargement). This data is consistent with a number of previous studies that showed GALC is expressed and is important in neurons^8, 32^. The unexpected distribution of GALC encouraged us to carefully determine the cellular expression of GALC in development. We therefore performed GALC colocalization experiments using various cell markers in the P5 brain. GALC protein was expressed in all brain cells including neurons, microglia, astrocytes, and OL lineage cells (Fig. 6d-e). The majority of GALC at P5 was expressed in neurons and was far higher than the expression in OL-lineage cells, astrocytes or microglia. Furthermore, neuronal GALC expression was developmentally influenced and peaked at P5 (Fig. 6f-g), similar to RNA and activity data (Fig. 6a-c). These results suggest a role for GALC in early neuronal development.

**Figure 6.**
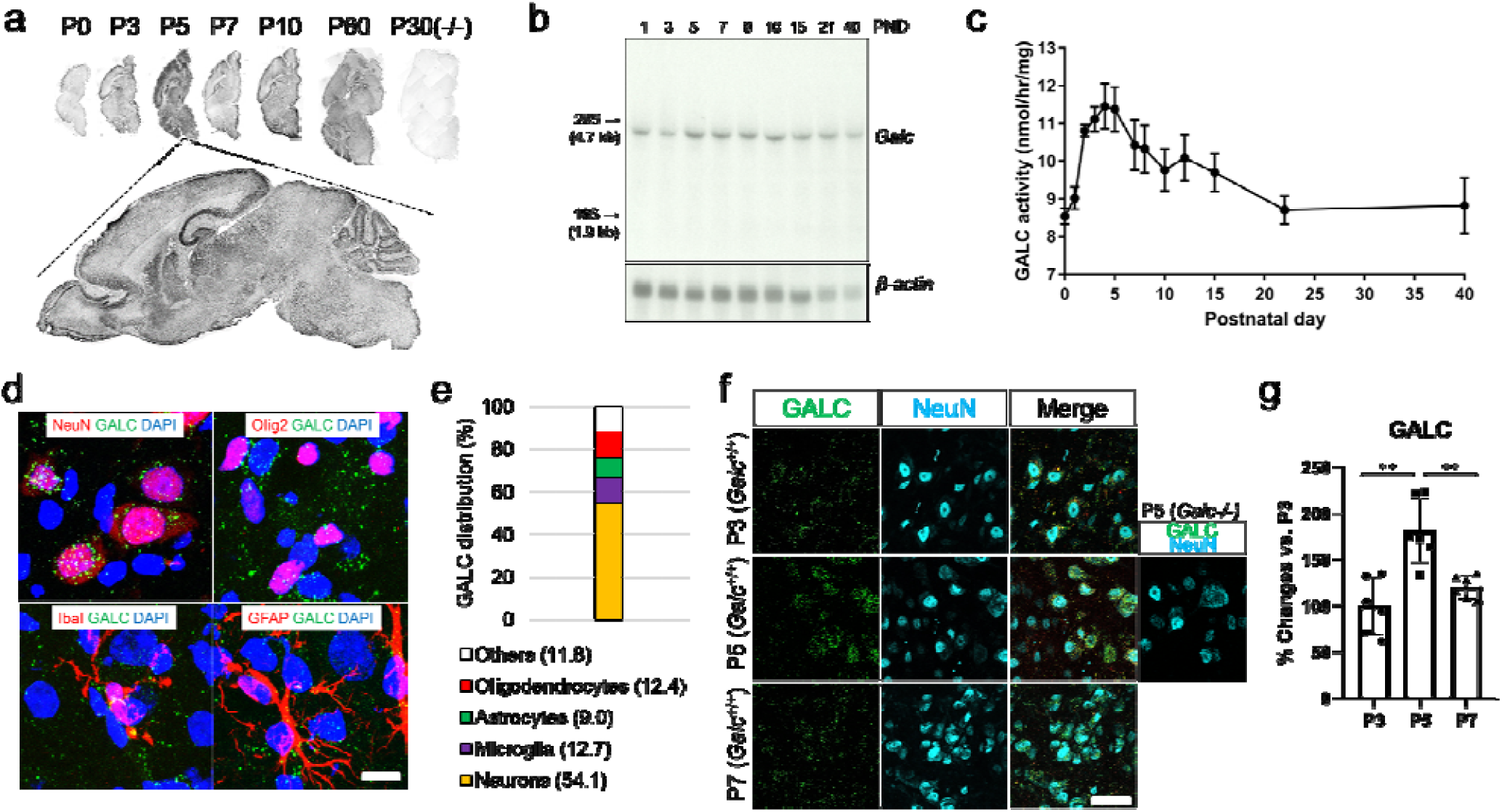
GALC levels reach their highest peak at P5 in the mouse brain. (**a**) *In situ* hybridization and (**b**) Northern blot analyses reveal that *Galc* transcript expression is low at P0, which then increase and reach their highest peak at P5 in WT brain. (**c**) GALC enzyme activity levels also show similar pattern of changes. (**d**) Immunohistochemistry on P5 brain shows that GALC protein is expressed in neurons (NeuN), microglia (IbaI), astrocytes (GFAP), and oligodendrocyte lineage cells (Olig2). Scale bar=20 *µ*m. (**e**) Quantification of GALC in each cell type reveals neurons express more than 50% of GALC in P5 brainstem. (**f, g**) Confocal images of brainstem neurons double-stained for GALC (green) and NeuN neuronal marker (cyan) in the P3-7 *Galc*^+/+^ brain shows that GALC is highly expressed in NeuN positive cells of the P5 brainstem. Scale bar=50 *µ*m.

### Brainstem development requires GALC expression

The parallel findings of increased GALC expression at P5, along with increased susceptibility to deletion of GALC prior to P5, suggests that a developmental process dependent on GALC occurs at P5. This finding is particulary surprising, as *twitcher* symptomatology does not begin until P21, and therefore suggests the possibility of an early developmental defect occuring prior to hallmark of KD progression. To monitor neuronal development, we used Thy1.1-YFP reporter mice that express YFP at high levels in a sparse population of motor, sensory, and some central neurons^33^. Thy1.1-YFP was crossed with *Galc*-KO mice or WT controls. Confocal analysis showed a dramatic decrease in the density of YFP-axons in the brainstem of P35 symptomatic *Galc*-KO; Thy1.1-YFP mice (Fig. 7a). Coronal sections confirmed that pyramidal, pontine retucular, trigeminal and gigantocellular nuclei were dramatically affected in the pons and medulla of *Galc-*KO; Thy1.1-YFP mice (Fig. 7b:”1”). Instead, the cerebellum and hippocampus were only moderately changed, while other brain rostral regions were not affected (Fig. 7b:”2, 3”). High power neuronal magnification further emphazied the region specific nature of pathology seen in the *Galc*-iKO mice (Fig. 7c-d), ultimately suggesting that hindbrain pathology is likely a major consequence of KD pathogenesis. Mutant axons sometimes appeared with swellings, breaks or transections, suggesting severe structural disruption with axonal degeneration (Fig. 7e). To determine if the reduction in Thy1.1-YFP signal reflected neuronal/axonal damage secondary to demyelination and inflammation, pre-symptomatic mice were analyzed at P14 (Fig. 7f). Intriguingly, a similar reduction of Thy1.1-YFP was seen in the brainstem of P14 *Galc*-KO mice, when clinical symptoms are normally absent. In line with previous studies, there was no evident microglial pathology at this early, ‘asymptomatic’ timepoint (Fig. 7g-h and Supplementary Fig. 2). Instead, we suspect that the reduced neuronal signal reflects a neural developmental perturbation in *Galc*-KO brains.

**Figure 7.**
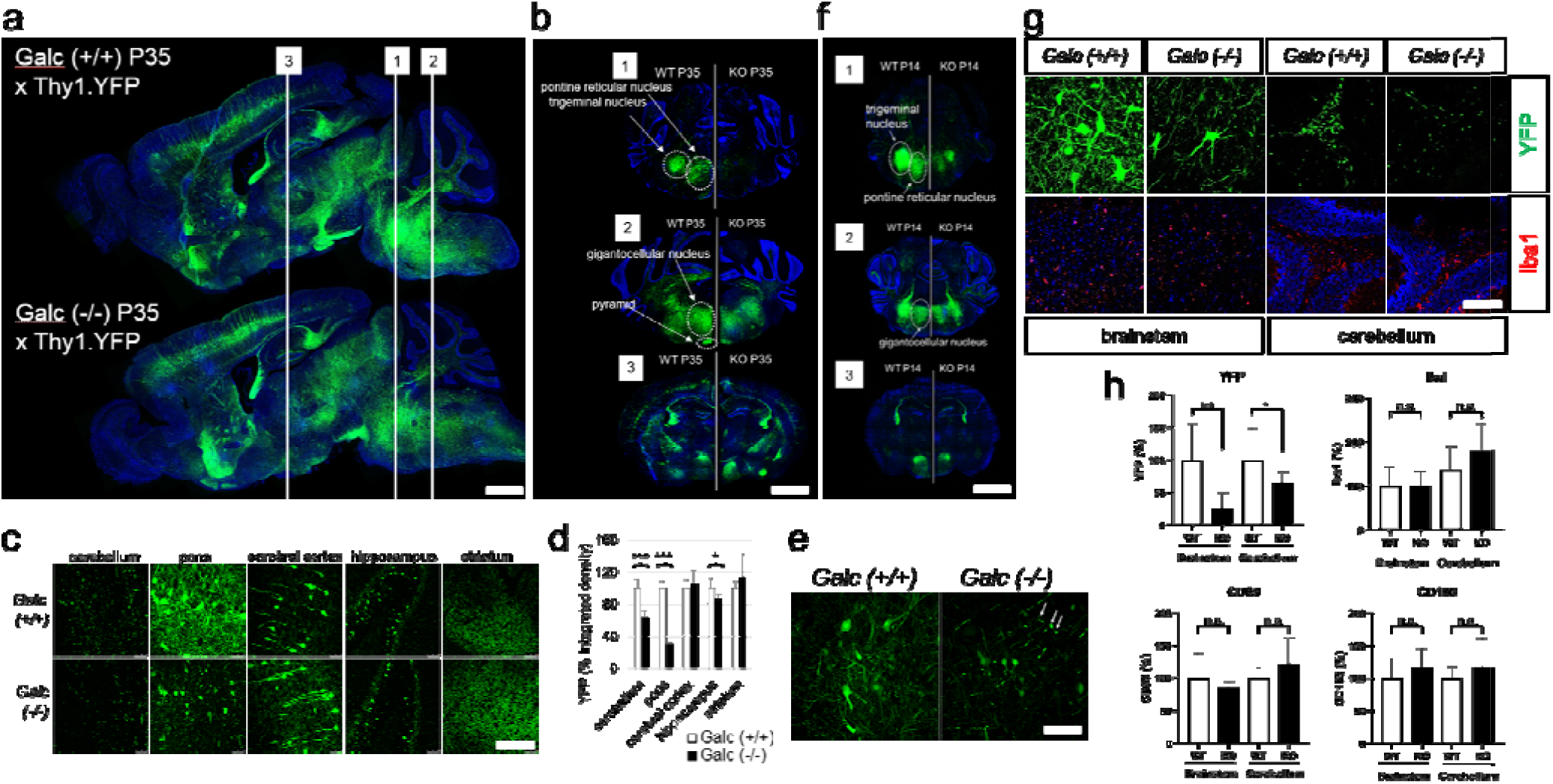
Brainstem is developmentally affected by GALC deficiency. (**a**) Double transgenic mice of P35 *Galc*-/-; Thy1.1-YFP shows that a decrease in the density of YFP-neurons/axons in the brainstem, compared to WT. (**b**) Coronal sections reveal pontine reticular/trigeminal/gigantocellular nuclei and pyramidal structures are dramatically affected (**c**) Highly magnified images and quantification (**d**) shows YFP signals significantly reduced in the cerebellum and pons (brainstem) and hippocampus. (**e**) Mutant axons in *Galc*-KO appeared with swellings (arrow), break or transections, indicating axonal degeneration. (**f**) P14 *Galc-/-*; Thy1.1-YFP brain has the same pattern of YFP reduction in trigeminal, pontine reticular, and gigantocellular nuclei, compared to WT. (**g**) Iba1-positive microglia are not activated in the brainstem and cerebellum of P14 *Galc*-KO, while YFP signals were much reduced. (**h**) Quantification of immunostained signals showed that microgliosis markers IbaI, CD68, and CD163 were not significantly changed, whereas YFP signals were much down-regulated. Scale bars: [**a**] 2 mm; [**b, f**] 150 *µ*m; [**c, e, g**] 50 *µ*m.

To determine if the attenuated neuronal signal was influenced by GALC expression in early development, we crossed the same Thy1.1-YFP mice with our *Galc*-iKO system. We then analyzed brains of P35 mice that were either induced at P3 (*Galc*-iKO≤P4) or P7-8 (*Galc*-iKO≥P6) (Fig. 8a). Comparing induction at both timepoints revealed that the brainstem, and in particular the pons, had a dramatic reduction of YFP-neurons/axons in the *Galc*-iKO≤P4, when compared to the *Galc*-iKO≥P6 (Fig. 8b-c). Other brain sub-regions were not affected as significantly. To confirm if this neuronal effect was the consequence of a developmental process, and not secondary to canonical KD pathogenesis, brains were analyzed immediately after 24 hours after the last tamoxifen administration (Fig. 8d). In line with a developmental defect, neuronal YFP signal in the brainstem was reduced immediately after tamoxifen induction in *Galc*-iKO≤P4 mice (Fig. 8e-f). Although a similar trend of reduction was also observed by temporal *Galc* deletion starting at P7 (*Galc*-iKO≥P6), this finding was not statistically significant (Fig. 8f). Taken together, these results suggest that GALC expression at P5 is critical in the development and stability of brainstem neurons.

**Figure 8.**
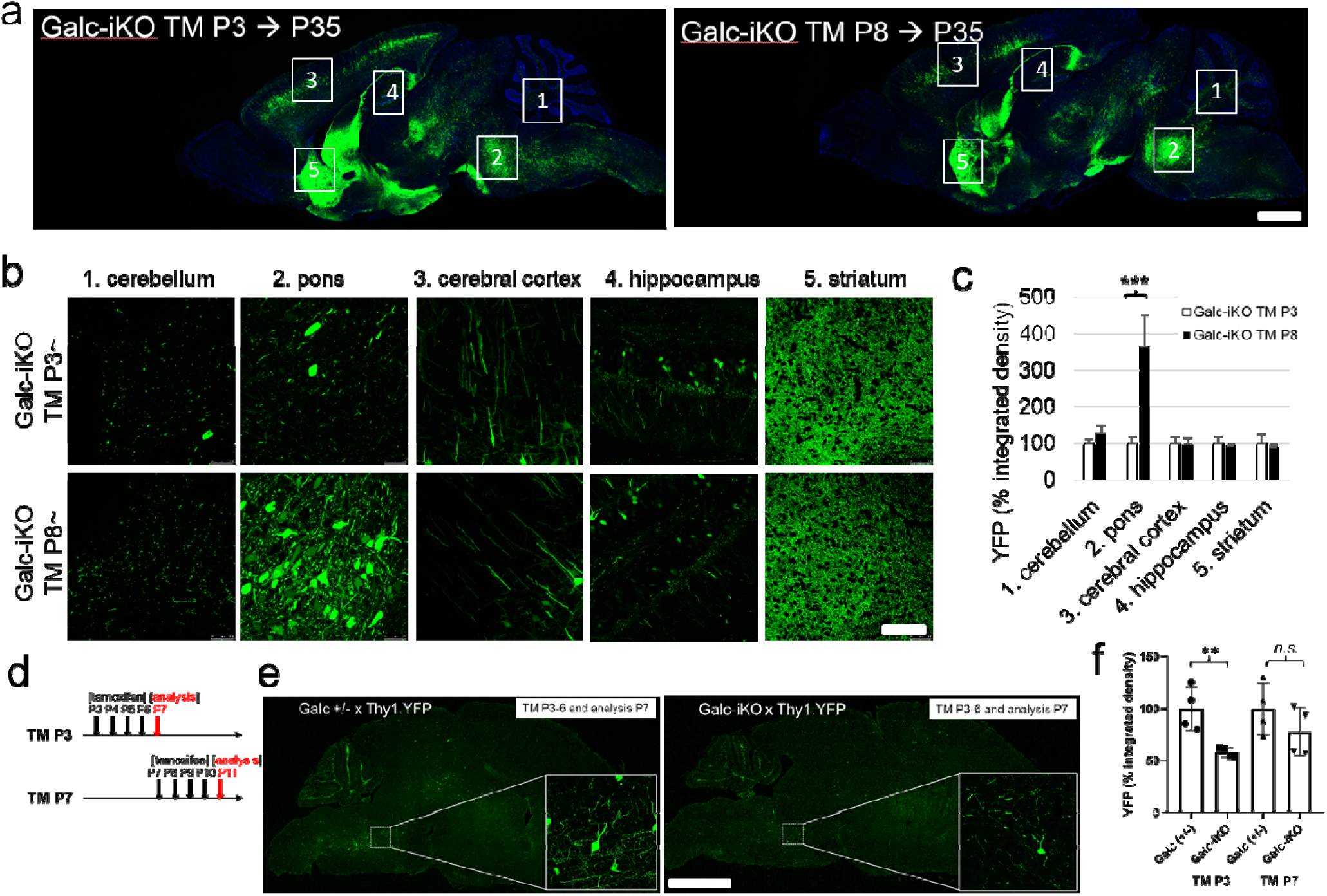
Brainstem development is affected during the critical period P4-6. (**a**) *Galc*-iKO;Thy1.1-YFP shows that a more decrease in the density of YFP-neurons/axons in the brainstem when induced at P3, compared to the induction at P8. Highly magnified images (**b**) and quantification of YFP density (**c**) shows a significant change of signals in the pons of *Galc*-iKO (TM P3) vs. iKO (TM P8). (**d-f**) Monitoring of neuron-specific YFP signals 24 hours after *Galc* deletion reveals that signification reduction of YFP intensity by induction at P3 but not by at P7. Scale bars: [**a, e**] 2 mm; [**b**] 50 *µ*m.

### *Galc* deficiency causes increase of a population of immature neurons

Although white matter and myelinating oligodendrocytes are thought to be the main contributors to Krabbe pathogenesis, they are not yet fully differentiated before P6. Furthermore, because neurons expressed the highest levels of GALC at P5 (Fig. 6), we explored the effect of GALC on neuronal development in the brainstem. T-brain-1 (TBR1), a brain-specific T-box transcription factor, plays a critical role in brain development. TBR1 expression is highest in immature neurons, at the embryonic stage of brain development, and is gradually reduced as neurons mature^34^. Interestingly, the number and overall intensity of TBR1 positive cells were significantly increased in the brainstem of *Galc*-KO mice compared to WT during P3-7 (Fig. 9a-c), suggesting GALC may be involved in the maturation step of neurons from an immature stage. The same analysis in *Galc*-iKO mice showed that the brainstem of *Galc*-iKO≤P4 had a dramatic increase of TBR1 positive cell bodies and intensity compared to controls, 24-hour after the induction (Fig. 9d-f). However, the TBR1 signals in the brainstem of *Galc*-iKO≥P6 did not show significant change, suggesting *Galc* deletion before P4 is critical for the persistent TBR1 expression.

**Figure 9.**
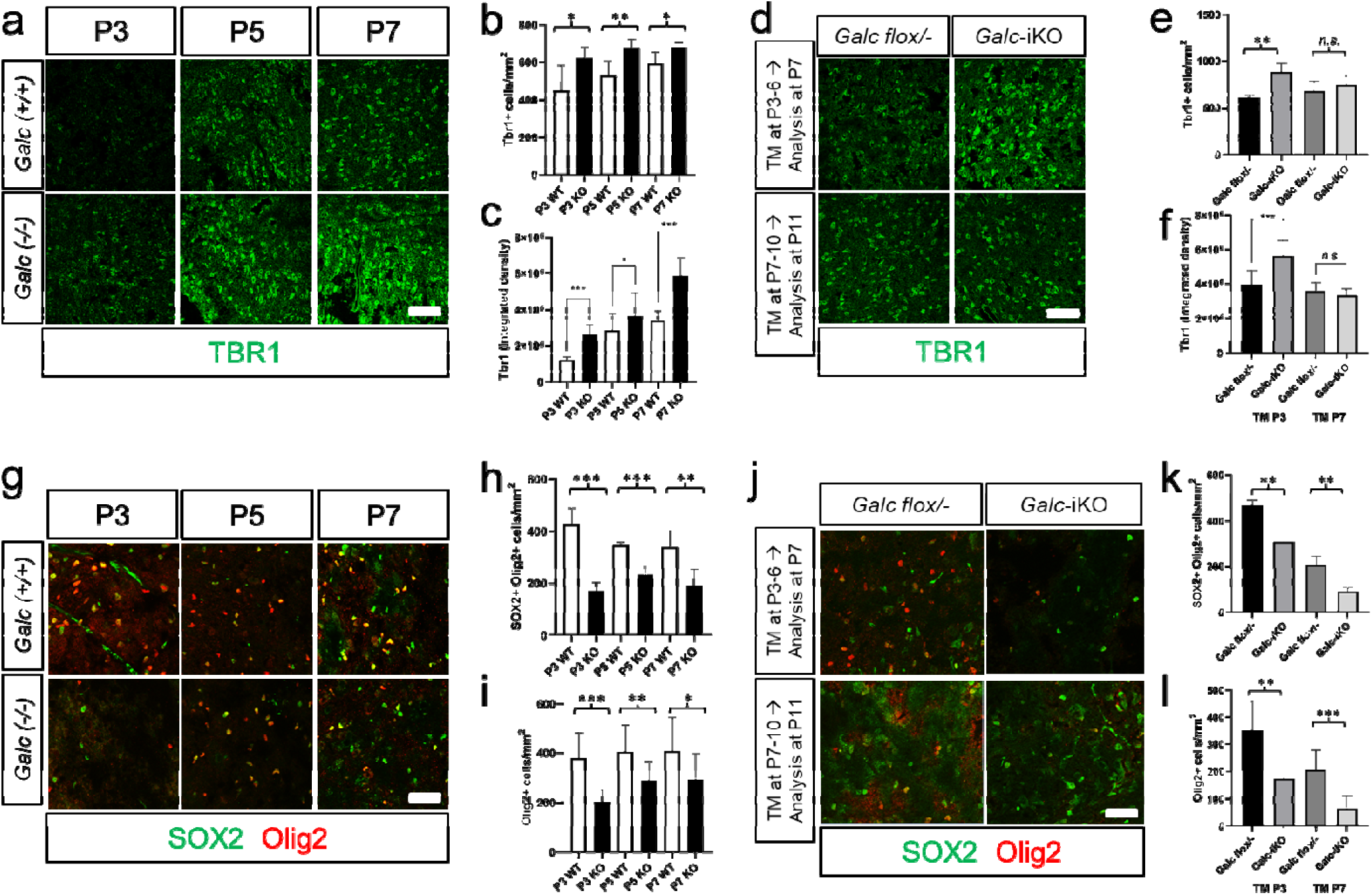
TBR1+ immature neurons are increased in the brainstem of *Galc* mutants, whereas SOX2+Olig2+ OPCs are reduced by GALC deficiency. (**a**) TBR1 positive cells were counted by immunohistochemistry on the brain sections of *Galc*-WT and KO at P3, P5 and P7. TBR1+ cells (**b**) and intensities (**c**) are significantly increased in *Galc*-KO in all ages. (**d**) Tamoxifen were induced starting at P3 or P7 in *Galc*-iKO, and after 24 hours TBR1-positive cells were stained. Quantification of TBR1+ cell numbers (**e**) and intensities (**f**) reveals *Galc* deletion at early time (P3) significantly increases TBR1+ cell number and intensity, however the change of *Galc*-iKO brain induced at P7 was not significant. (**g-i**) SOX2 and Olig2 positive cells were counted by immunohistochemistry on the brain sections of *Galc*-WT and KO at P3, P5 and P7. In all three ages, the numbers of SOX2+Olig2+ and Olig2+ cells were reduced by *Galc-*KO. (**j-l**) Tamoxifen were administrated starting at P3 or P7 in *Galc*-iKO, and after 24 hours SOX2 and Olig2 positive cells were counted. Both inductions significantly reduce the population of both SOX2+Olig2+ and Olig2+ cells, suggesting oligodendrocytes are affected by *Galc* deletion but is independent of the critical period P4-6. Scale bar=50 *µ*m.

The neuronal marker NeuN begins to be expressed during early embryogenesis in postmitotic neuroblasts and remains expressed in differentiating and terminally differentiated neurons thereafter^35^. The overall NeuN positive cell number was not different between WT and KO brainstems at P3, 5, and 7 (Supplementary Fig. 3a, b). This is probably due to the fact that NeuN detects both mature neurons and undifferentiated neuroepithelial cells. We also measured proliferating neurons by EDU labeling for 24-hour prior analysis. EDU positive neurons were very rare during the period and were not different between WT and KO (Supplementary Fig. 3a, c). Since EDU labels cells only during the S phase of cell cycle, we also measured Ki67 signals which are expressed during all stages of cell proliferation including the G1, S, G2, and M phases of cell cycle. Ki67 labeled cell number was also not changed in *Galc*-KO (Supplementary Fig. 3d), suggesting proliferation of neurons is not affected by *Galc* deficiency postnatally. Despite no change of neuronal cell number in early developmental time, there was significant reduction in P40 *Galc*-KO brainstem when the mouse is moribund (Supplementary Fig. 3e) presumably by apoptosis (Supplementary Fig. 3f, g), though it is not clear if this reduction is due to neuron-autonomous or secondary effects of other cell’s pathogenesis.

### SOX2+Olig2+ OPCs are reduced in the brainstem by *Galc* deficiency

During the critical period P4-6, GALC is expressed not only in neurons but also in other cell types including OL-lineage cells (Fig. 6d, e). The murine pons, which is a core structure of brainstem, quadruples in volume shortly after birth and peaks at P4, preceding myelination^36^. During this period, postnatal SOX2+Olig2+ OL-progenitor cells (OPC) expand 10-18 fold into the OL-lineage cells that later comprise the more than 90% of adult pons OLs. Therefore, in addition to neurons, it is possible that OPCs could be a key player affecting the development of KD brainstem during P4-6. To test this hypothesis, we counted OPC numbers in the brainstem of *Galc*-KO and WT at P3, P5 and P7 by immunostaining with the OL-lineage marker Olig2 and neural stem cell lineage marker SOX2. Interestingly, both numbers of SOX2+/Olig2+ and Olig2+ cells were dramatically reduced in the brainstem of the *Galc*-KO, compared to WT at all P3, 5, and 7 (Fig. 9g-i and Supplementary Fig. 4), suggesting that GALC has a specific role in the development of OLs in the postnatal brainstem. As before, we analyzed the number of OPCs in the brains of both *Galc*-iKO≤P4 and *Galc*-iKO≥P6. Interestingly, within 24 hours of tamoxifen injection, SOX2+/Olig2+ and Olig2+ cell numbers were dramatically reduced in the brainstems of both *Galc*-iKO≤P4 and *Galc*-iKO≥P6 (Fig. 9j-l and Supplementary Fig. 5). These data suggest that GALC expression is required for the expansion of the OPC population during brainstem development, but is independent of the critical period P4-6.

### Neuron-specific perinatal *Galc* influences neuronal differentiation without affecting OPCs

Analysis of *Galc*-iKO highlights the importance of GALC in neurons during early brainstem development. To determine if neuron-specific GALC ablation is sufficient to attenuate the maturation of neurons, we used pan-neuron specific Thy1-Cre/ER^T2^ mice^37^ to induce neuron-specific temporal *Galc*-CKO mice. Although the Thy1-Cre/ER^T2^ line is well characterized in the literature, we first confirmed Cre specificity and efficiency. To test both, tamoxifen was injected into Thy1-Cre/ER^T2^;tdTomato starting at P2 tamoxifen at a dose of 25 μg per gram of bodyweight for four consecutive days. At P10 and P30, the animals were analyzed for the specificity of Cre expression and efficiency of Cre-loxP recombination. The tdTomato protein is efficiently expressed in Thy1 positive cells in whole brain including cerebellum, brainstem, cerebral cortex and corpus callosum (Supplementary Fig. 6). 60-80% of Thy1 signals colocalized with tdTomato in the brain at both P10 and P30, indicating substantial specific recombination in neurons. Other cell markers Olig2 (oligodendrocytes), GFAP (astrocytes), and Iba1 (microglia) were minimally co-labeled with tdTomato, suggesting that Thy1 promoter driven Cre expression is specific to neurons. Next, we analyzed the population of immature neurons in the brainstem of Thy1-Cre/ER^T2^ driven neuron specific *Galc*-CKO induced starting at P2. As before, the analysis of mice at P60 showed increase of TBR1 cell number and intensity in the brainstem of neuron specific conditional mice vs WT (Fig. 10a-c). Similar to *Galc*-KO (Supplementary Fig. 3a, b), the overall NeuN positive cell number was not different between WT and CKO brainstems at P60 (Fig. 10e). However, the number of Olig2-positive cells in the brainstem was not changed (Fig. 10d, f). This data suggest that neuronal GALC affects the maturation of neurons in a cell autonomous manner, but not the expansion of the OPC population.

**Figure 10.**
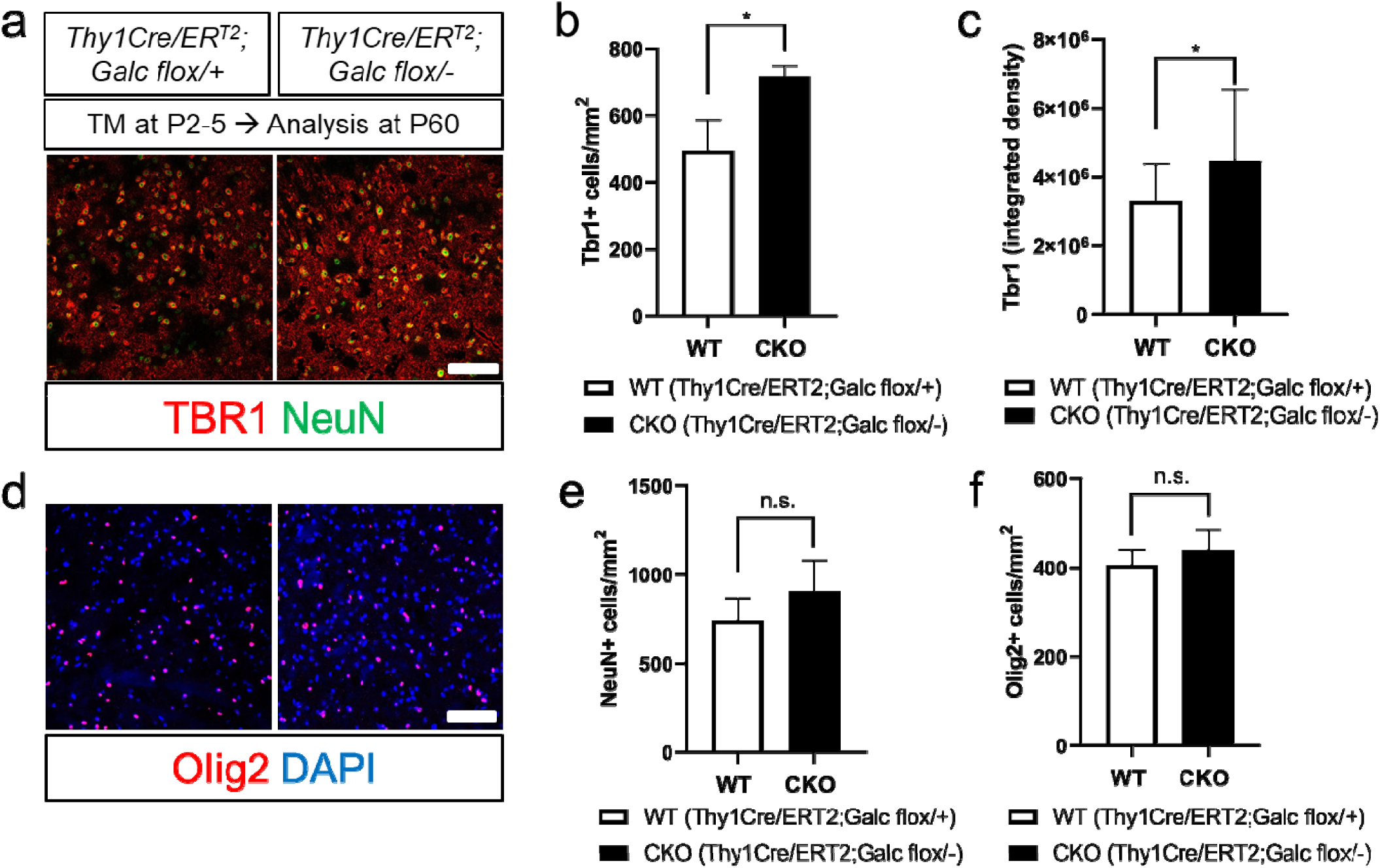
TBR1+ immature neurons are increased in the brainstem of neuron-specific *Galc* knockouts. Immunostaining of TBR1 on the brainstem of Thy1-Cre/ER^T2^ driven *Galc* deletion induced starting at P2 (**a**) showed that the number of TBR1+ cells (**b**) and intensities (**c**) are significantly increased in Thy1-Cre/ER^T2^; *Galc*^flox/-^ at P60. However, there were no changes in the numbers of NeuN (**e**) and Olig2+ cells (**d, f**). Scale bar=50 *µ*m.

## Discussion

### Temporal Requirement for GALC in KD pathogenesis

To better understand the role of GALC in development, we used a conditional mutagenic approach and generated a novel mouse model of KD. We were particularly interested in the role of early postnatal GALC function as empirical clinical evidence suggests that gene and stem cell therapy is more efficacious if delivered in a poorly defined pre-symptomatic window^38^. Our approach was to methodically induce *Galc* deletion at various postnatal timepoints, thereby determining key postnatal events impacted by GALC deficiency. While we expected GALC to be important at a time-period that coincided with myelination, we were surprised to find that GALC deficiency abruptly accelerated mortality when ablated before P4 compared to after P6 in C57BL6 mice. The protective role of perinatal GALC is particularly surprising and suggestive of a novel function unrelated to its canonical role in myelination.

This developmental role for GALC likely correlates with the observed clinical benefit of early, pre-symptomatic KD treatment. For example, pre-symptomatic infants transplanted before one month old have better outcomes than older pre-symptomatic infants^38^. KD pathogenesis may begin very early in development, perhaps even prenatally. While the time scale between mice and humans is considerably different, the sequence of key events in brain maturation such as neurogenesis, synaptogenesis, gliogenesis, and myelination between the two is consistent^39^. It was estimated that the mouse CNS at P1–3 corresponds to a gestational age of 23–32 weeks in humans, P7 to 32–36 weeks and P10 to a term infant, at least in regards to white matter development^40^. Therefore, we anticipate that if our hypothesis on the early critical period of vulnerability is correct, then in-utero treatments should have better outcomes than conventional postnatal treatment.

We also explored the possibility that the worsened clinical phenotype seen in early induced mice reflected variable postnatal recombination. We were particularly concerned that the developing blood-brain barrier (BBB) in early neonatal mice would cause differences in tamoxifen uptake. Instead, we found recombination was similar among our variable induction timepoints (Fig. 3). This data fits with the fact that tight junctions between cerebral endothelial cells (the morphological basis for BBB impermeability) are functionally effective as soon as the first blood vessels penetrate the parenchyma of the developing brain^41^, and also evidenced by analyzing tamoxifen metabolites in the postnatal brain^24^. Nonetheless, it is worth noting that the earliest induced *Galc*-iKO mice, occurring on postnatal day P0, still had significantly longer life-spans (∼P60) than either *Galc*-KO or *twitcher* (∼P45) (Fig. 1). This may indicate that the inducible Cre-loxP system is incomplete and thereby delays the KD phenotype^42^. Alternatively, deletion of *Galc* prior to birth may be required to fully recapitulate the global KD phenotype.

Another interesting and unexpected finding of our study was that residual GALC returned in the late course of the disease models (Fig. 3h). Our inducible Cre-LoxP system is dependent upon tamoxifen expression for continued recombination events, thus it seems plausible that substantial GALC activities could eventually return after *Galc* ablation, especially if a non-recombined minority of cells repopulated over time. Intriguingly, despite the substantial GALC activity that returned in all *Galc*-iKO mice, the disease course remained unaltered and mice continued to deteriorate and inevitably reached moribundity (Fig. 2b, c). This finding may correlate with recent gene therapy trials for a number of LSDs in which increased enzyme activity was detected at the whole parenchymal brain level, but unfortunately resulted in minimal clinical improvements^43, 44^. This suggests that providing GALC in specific developmental period is critical to treat KD.

### Cell and Region-specific Requirement for GALC in Brain Development

After elucidating a developmental critical period that requires GALC, our next task was to discern the cellular, regional and functional mechanism by which GALC is protective. Our work illustrates that GALC is not only required by myelinating cells, but is also expressed in many other brain cells as shown in Figure 6. At the perinatal timepoint, GALC is most highly expressed in neurons, suggesting a novel neuron autonomous role for GALC in early postnatal development. It is also known that GALC in neural stem cells maintains a subventricular zone neurogenic niche during early postnatal period^45^. While the majority of neurons are generated embryonically, many neuronal populations must mature and differentiate postnatally. For example, the dentate gyrus (DG) is marked by substantial postnatal neuronal maturation, with a high number of immature granule cells (GCs) present during in the first two weeks of postnatal life. These GC neurons then progress toward more mature patterns over the next few weeks of DG maturation^46^. Similarly, postnatal refinement of neurons of the visual cortex is critical for the intrinsic properties and plasticity needed for proper function and network activity^47^. Furthermore, most pontine circuits are postnatally acquired or refined^36^, which contains nuclei that relay signals from the forebrain to the cerebellum, along with nuclei that deal primarily with vital functions^48^. Therefore, our finding that GALC influences postnatal neuronal differentiation, while surprising, is not unprecedented.

We show that brainstem development is dependent on intrinsic GALC expression by neurons. Specifically, brainstem neurons lacking GALC had perturbed differentiation and were in a more immature state. Neurons of this region ultimately had more axonal atrophy and degeneration when *Galc* was deleted between P4-6. These pathological findings were not equal among all brain regions and were most pronounced in neurons of the brainstem. Interestingly, defective neurogenesis of the hindbrain and midbrain regions were previously noted in a zebrafish model of KD^49^. The brainstem was also previously implicated in KD as the region where pathology first develops^50^. While the functional consequence of brainstem dysfunction is difficult to directly assess, it is known to serve as an important relay between the forebrain, cerebellum and other nuclei associated with vital functions^48^. In fact, most children who develop KD in infancy die before the age of 2, often from respiratory failure^51^ and suspected autonomic dysfunction. In addition, even pre-symptomatic KD infants treated with HSCT continue to develop substantial motor impairments because of brainstem dysfunction involving corticospinal tract pathology^27^. Therefore, we suspect that GALC-dependent brainstem dysfunction may directly influence KD pathogenesis.

Despite our conclusion that P5 GALC influences brainstem neuronal differentiation, the mechanism by which this happens remains unknown. Prior to the onset of overt clinical symptoms, neuronal swellings, varicosities and transected fibers were already detectable in *Galc-*deleted mutants (Fig. 7f-h, 8d-f). These findings preceded overt neuronal loss, as reductions in neuronal numbers only occurred shortly before death (Supplementary Fig. 3). Similarly, previous reports in the *twitcher* mouse emphasized that axonal abnormalities were evident very early in postnatal development (P7), while neuronal apoptosis was detectable only after disease onset (after P30). This suggests that axonal stress and dysfunction precedes overt neuronal apoptosis in *twitcher*^8^, a process which is consistent with the pathophysiology of other neurological disorders^52^. Since lysosomes are the essential building blocks for synaptic biogenesis^53^, it is conceivable that the degenerative process may be related to neuronal synaptic function at the axon terminal. Here, structural and functional defects may begin to impact synaptic efficiency, thereby causing overt neuronal toxicity^54^, apoptosis and ultimately overall brain development^55, 56^. We suspect that immature brainstem neurons, influenced by autonomous GALC deficiency, are more vulnerable to toxins like psychosine while also being less able to respond to local environmental signals, and are eventually eliminated^57^.

In addition to neuronal differentiation abnormalities, we observed that OL-lineage cell expansion was significantly reduced upon perinatal *Galc* deletion in the brainstem perinatally. Due to the common origin of neurons and OL-lineage cells, namely neural stem cells, it seemed conceivable that a dynamic interplay between the differentiation of OLs and neurons influenced proper brainstem development. However, our study showed that neuronal GALC did not influence the expansion of the OPC population during the critical period P4-6 (Fig. 10), suggesting a neuron-autonomous role of GALC for neuronal maturation exclusively. In fact, although myelination of the central nervous system appears normal in the early postnatal life of the *twitcher* mouse^14^, axonal swellings and varicosities occur as early as P7^8, 14^. These early axonal phenotypes, in the absence of demyelination, suggest a neuron autonomous effect of GALC deficiency. Also we cannot exclude the possibility that GALC may directly regulate the activation of TBR1 in the progression of neuronal maturation. Furthermore, the absence of GALC in mutant neurons results in a three-fold increase of psychosine, which may be sufficient to induce morphological changes in mutant neurons^8^.

In summary, our findings highlight a previously unidentified critical period in which GALC is required for neuronal brainstem development. The sudden and abrupt change in the clinical phenotype of *Galc*-iKO mice induced before or after P5 suggests a highly dynamic developmental process related to the function of GALC. This finding is particularly interesting as P0-P10 represents a pre-symptomatic timepoint in *Galc*-KO mice, occurring well before the majority of myelination in the murine brain. These findings are novel and surprising, but may correlate with the pre-symptomatic therapeutic window seen in treating KD patients. Further studies are required to elaborate on the specific cellular mechanisms which require GALC for brainstem development and if similar processes occur in other LSDs.

## Methods

### Generation of a conditional *Galc* floxed allele mouse

A *Galc* targeting vector (Supplementary Fig. 1a) from a BAC clone harboring a mouse *Galc* gene (bMQ-165I15, Source Bioscience, UK) was made by using a genetic recombineering technique (GeneBridges GmbH, Germany). Of the 17 *Galc* exons, we decided to flank exon 9 by two loxP sites, because the region from exons 7-10 on the *Galc* gene is consistently expressed among all splice variants (Ensembl Genomes). Furthermore, removal of exon 9 leads to a frame shift mutation in the remaining protein coding transcript. The targeting vector was then injected into embryonic stem (ES) cells from 129S6 mice, ultimately yielding 225 surviving clones. PCR screening of the vector’s short arm junction led to the selection of 7 *Galc*-targeted positive clones, which were confirmed further by Southern blot (Supplementary Fig. 1b) and DNA sequencing of both arm-junctions. Two positive ES cells were injected into blastocysts, and both cells generated chimeras successfully. One chimera transgene was germline transmitted (**r** in Supplementary Fig. 1a), which was crossed with Del-FLPe mice (JAX#009086)^58^ to remove the FRT-neomycin-FRT cassette. The resultant floxed mice (**f** in Supplementary Fig. 1a) were mated to constitutive ubiquitous Cre (CMV-Cre; JAX#006054)^19^ to validate if *Galc* can be completely deleted by Cre-loxP recombination (**-** in Supplementary Fig. 1a), and generate a null phenotype that would be comparable to *twitcher*. For genotyping of *Galc* floxed mouse, three primers were used; Primer-1 = 5’-CATCATCCTGTTTCCACAGG-3’, Primer-2 = 5’-AATATGTAGGGAGAGAGTGGTC-3’, Primer-3 = 5’-CTATTTTAAGGGAGTTCTGCCAGTG-3’. WT is 250 bp, loxP-floxed is 350 bp, and null allele is 450 bp.

### Animals

Experiments were conducted according to the protocols approved by the Institutional Animal Care and Use Committee of University at Buffalo and Roswell Park Cancer Institute. All animals were maintained on the congenic background of C57BL/6N. Breeder C57BL/6N mice were purchased from Charles River (Wilmington, MA). CAG-Cre/ER^T^(JAX#004682), CMV-Cre (JAX#006054), tdTomato (JAX#007905), Thy1.1-YFP (JAX#003782), and Thy1-Cre/ER^T2^(JAX#012708) were purchased from The Jackson laboratory (Bar Harbor, ME). For Cre/ER^T^-mediated recombination, a 5 mg/ml Tamoxifen (Sigma-Aldrich) solution was prepared in autoclaved corn oil (Sigma-Aldrich). To achieve efficient *Galc* recombination by tamoxifen, multiple pilot experiments were conducted with varying doses (25-100 μg per gram of body weight) and times (2-5 consecutive days) of tamoxifen injection. Finally, perinatal CAG-Cre/ER^T^; *Galc*-floxed mice were injected intraperitoneally for 4 consecutive days (total 4X) with 25 *µ*g/gram body weight, in pups between P0-P10 (data not shown). This was the maximum achievable dosage while avoiding tamoxifen-induced gastric toxicity^59^. It has been reported that tamoxifen affects glucose/lipid metabolism^60^ and myelination^61^, which could theoretically affect the survival of GALC-deficient mice. To exclude this potential confounding factor, we treated *Galc*-KO mice with tamoxifen using the same paradigm, starting injections at P4. Importantly, tamoxifen did not affect the life span of *Galc*-KO mice (40-45 days, n=3), arguing against the possibility of confounding tamoxifen toxicity. For proliferation assays, mice were pulsed with 100 *µ*g per gram of body weight of 5-Ethynyl-2’-deoxyuridine (EdU) (Sigma-Aldrich) at 24 hours before sacrifice by injection intraperitoneally.

### Tissue and Immunohistochemistry

Mice at defined ages were anesthetized, sacrificed and then perfused with ice-cold phosphate-buffered saline (PBS) followed by 4% paraformaldehyde (PFA). Brains and spinal cords were dissected, post-fixed in 4% PFA overnight, dehydrated in 30% sucrose at 4°C, embedded in OCT (Leica) and processed as cryosections with a thickness of 15 *µ*m. For immunohistochemistry, cryosections were permeabilized and blocked in blocking buffer (0.1% Triton X-100, 20% fetal bovine serum, 2% bovine-serum albumin in PBS) for 1 hour at room temperature and overlaid with primary antibodies overnight at 4°C. After washing with PBS, sections were incubated with fluorophore conjugated-secondary IgGs (Jackson laboratories). After washing (3X for 5 minutes) with PBS, coverslips were mounted with Vectashield (Vector Laboratories) mounting medium and DAPI. Primary antibodies used were GALC^62^, NeuN (EMD Millipore), Olig2 (Peprotech), GFAP (Sigma-Aldrich), Iba1 (Wako), CD68 (Bio-Rad), CD163 (Bio-Rad), tdTomato (Origene), SOX2 (R&D systems), TBR1 (Abcam), cleaved caspase-3 (Cell Signaling Technology), and Ki67 (Invitrogen). For colorimetric cleaved caspase-3 staining, SignalStain® Boost IHC Detection Reagent (Cell Signaling Technology) was used with Hematoxylin counterstain. Images were acquired and analyses performed whilst blinded to genotype. Area of fluorescence and fluorescence intensity were quantified with ImageJ (NIH).

### Cell counts and z-stacks

The cell type composition was assessed in matched sagittal brain sections (n>3 mice) selected within the region containing each cell specific markers (NeuN for neurons; GFAP for astroglia; Olig2 for oligodendroglia; Iba1 for microglia; TBR1 for immature neurons) and nuclear counterstaining (DAPI) in immunofluorescence followed by Leica SP5 confocal microscopic analysis. z-stacks were recorded utilizing sequential confocal images that were collected at 1 μm intervals covering 30 μm depth. Numbers for each cell type were acquired by manually counting cells.

### Transmission Electron Microscopy (EM)

Mice were first anesthetized with 250 mg/kg body weight avertin and then perfused with PBS and 2.5% gluta-aldehyde in phosphate-buffer and incubated in fixative for one week. After being post-fixed, spinal cords were dissected and embedded in Epon. Ultra-thin sections were cut and stained with uranyl acetate and lead citrate and then collected on grids. The pictures were taken with a Tecnai electron microscope.

### Western blot analyses

After homogenizing whole brains in RIPA buffer containing protease inhibitors (Roche) and PMSF, total protein extracts were separated by SDS-PAGE, transferred to PVDF membrane (Millipore), and blocked with 5% Skim milk or BSA in TBS-Tween20. Primary antibodies used were GAPDH (Chemicon), β-tubulin (Novus Biologicals), tdTomato (Origene), TLR2 (R&D systems), GFAP and HA (all from Sigma-Aldrich). Specific protein bands were quantified by means of ImageJ and Image Studio (LI-COR Biosciences), and the values (in pixels) obtained were normalized on those of the corresponding β-tubulin bands. Normalized values were then expressed as the percentage of values obtained from region-matched bands of control WT tissues.

### GALC enzyme assay

GALC activity was determined by the previously described method^2^. Briefly, snap frozen whole brains were homogenized in 10 mM sodium phosphate buffer, pH 6.0, with 0.1% (v/v) Nonidet NP-40 by using Dounce homogenizer. 1 μg of total brain lysates were mixed with 4-methylumbelliferone-β-galactopyranoside (final 1.5 mM) resuspended in 0.1/0.2 M citrate/phosphate buffer, pH 4.0, and AgNO_3_ (final 55 *µ*M) at 37°C for 1 hour. The enzymatic reactions were stopped by adding 0.2 M glycine/NaOH, pH 10.6. Fluorescence of liberated 4-ethylumbelliferone was measured on a spectrofluorometer (λex 360 nm; λem 446 nm).

### Northern-blot hybridization

Northern-blot analyses with 20 μg of total RNAs from mouse brains were performed as described previously^63^. The probes were prepared as the [α-^32^P] dCTP *Galc* exon fragment excised from the plasmid that harbors PCR product for exon 5-9 of mouse *Galc* (540 bp) and the mouse glyceraldehyde-3-phosphate dehydrogenase (508 bp).

### *In situ* hybridization

*Galc in situ* hybridization on cryosections was performed as described previously^64^. Briefly, cryosections of brain were incubated with digoxigenin (DIG)-labeled antisense riboprobes for murine *Galc*. The probe was synthesized using T3 RNA polymerase (Promega) and labeled with DIG RNA label mix (Roche). An anti-DIG antibody conjugated with alkaline phosphatase (Roche) was used to probe sections, which were stained with 5-bromo-4-cloro-3-indlyl phosphate/nitro blue tetrazolium (Roche) chromogenic substrates.

### Measurement of psychosine

Brain lysates were extracted in chloroform:methanol and partially purified on a strong cation exchanger column. After evaporation to dryness, each residue was dissolved in methanol and analyzed using LC-MS/MS^8^.

### Statistical analyses

Data collection and analysis were performed blind to the conditions of the experiments. Two tailed unpaired Student t-test with Bonferroni corrected and One way ANOVA were used for the differences among multiple groups according to the number of samples, except *G-ratio* analysis in which Welch’s t-test was used. Values of *P* < 0.05 were considered to represent a significant difference. Statistical tests were run in GraphPad Prism. Data are presented as mean ± SEM or SD.

## Acknowledgements

We would like to thank Aimee Stablewski in Roswell Park transgenic animal facility for the generation of *Galc* conditional allele mouse, Christopher B. Eckman in Biomedical Research Institute of New Jersey for providing the polyclonal GALC antibody, and Ed Hurley in the Hunter James Kelly Research Institute for technical support of confocal and electron microscopic analyses. The authors declare no competing financial interests.

## Supplementary Figures

**Figure S1.**
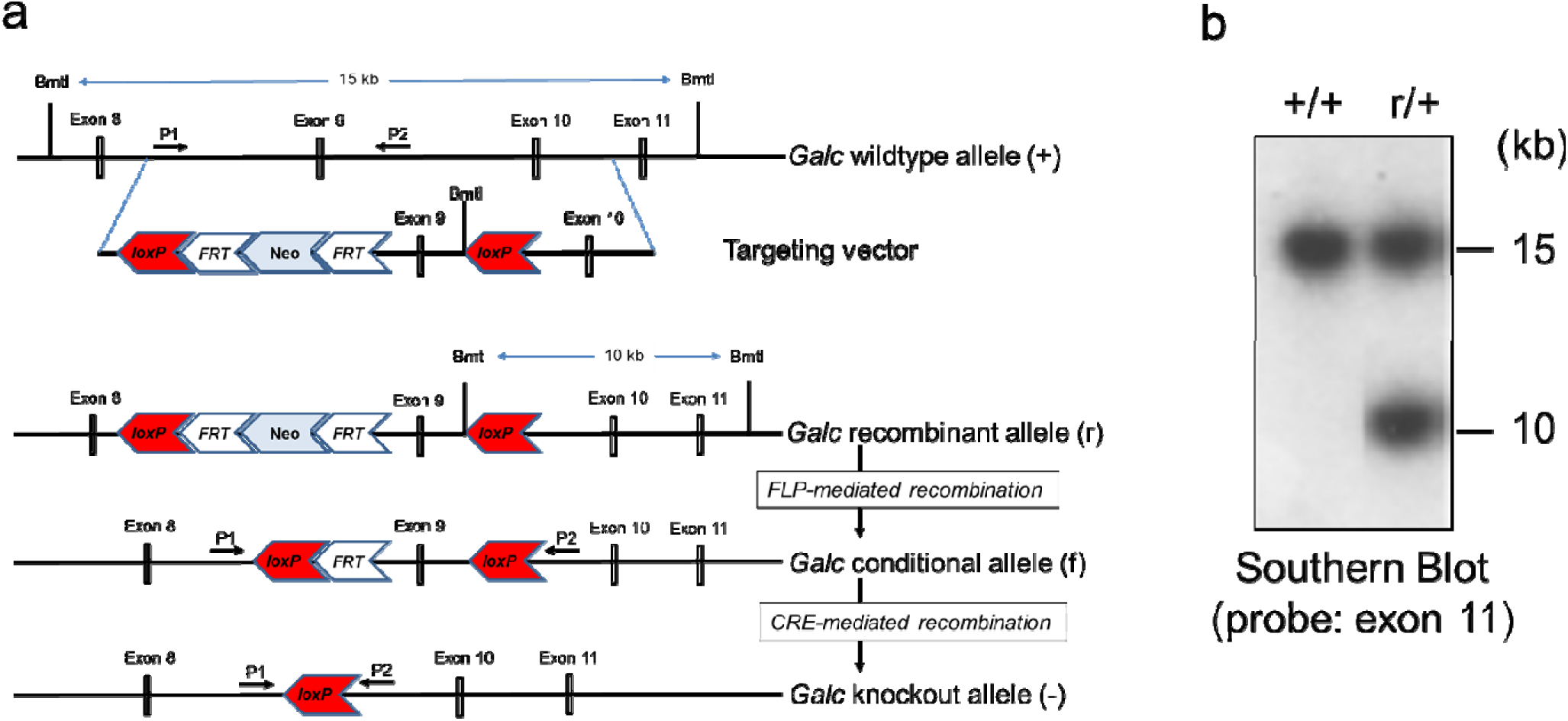
Design of conditional *Galc* floxed allele mouse. (**a**) Structure of the wild-type (+) locus, the targeting vector, the targeted recombinant allele (r), the conditional allele (f), and the deleted KO (-) *Galc* (exon 9) allele, the loxP and FRT recombination sites (red and empty, respectively), the *Bmt*I restriction enzyme sites and exons, and the P1-P2 primers for PCR genotyping are indicated. (**b**) Southern blot analysis of the wild-type (+/+) ES cells and of the recombinant (r/+) clones. *Bmt*I-restricted genomic DNA yielded 15- and 10-kb bands for the wild-stype and recombinant alleles, respectively, with the probe on exon 11 of the *Galc* gene.

**Figure S2.**
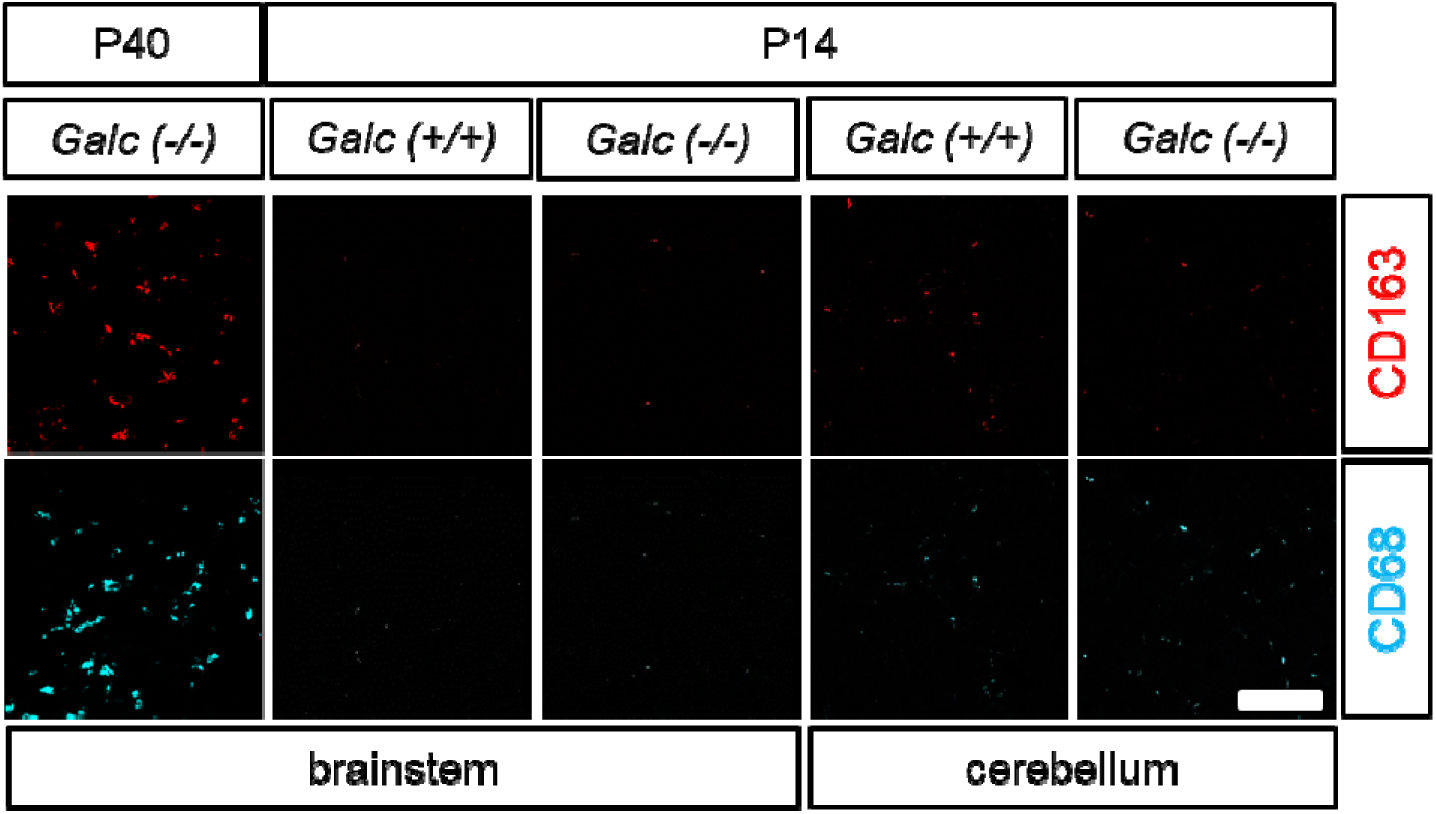
CD68 (red) and CD163 (cyan) positive microglia are not activated in the brainstem and cerebellum of P14 *Galc-*KO, which are highly increased in the moribund P40 KO brains. Scale bar=50 *µ*m.

**Figure S3.**
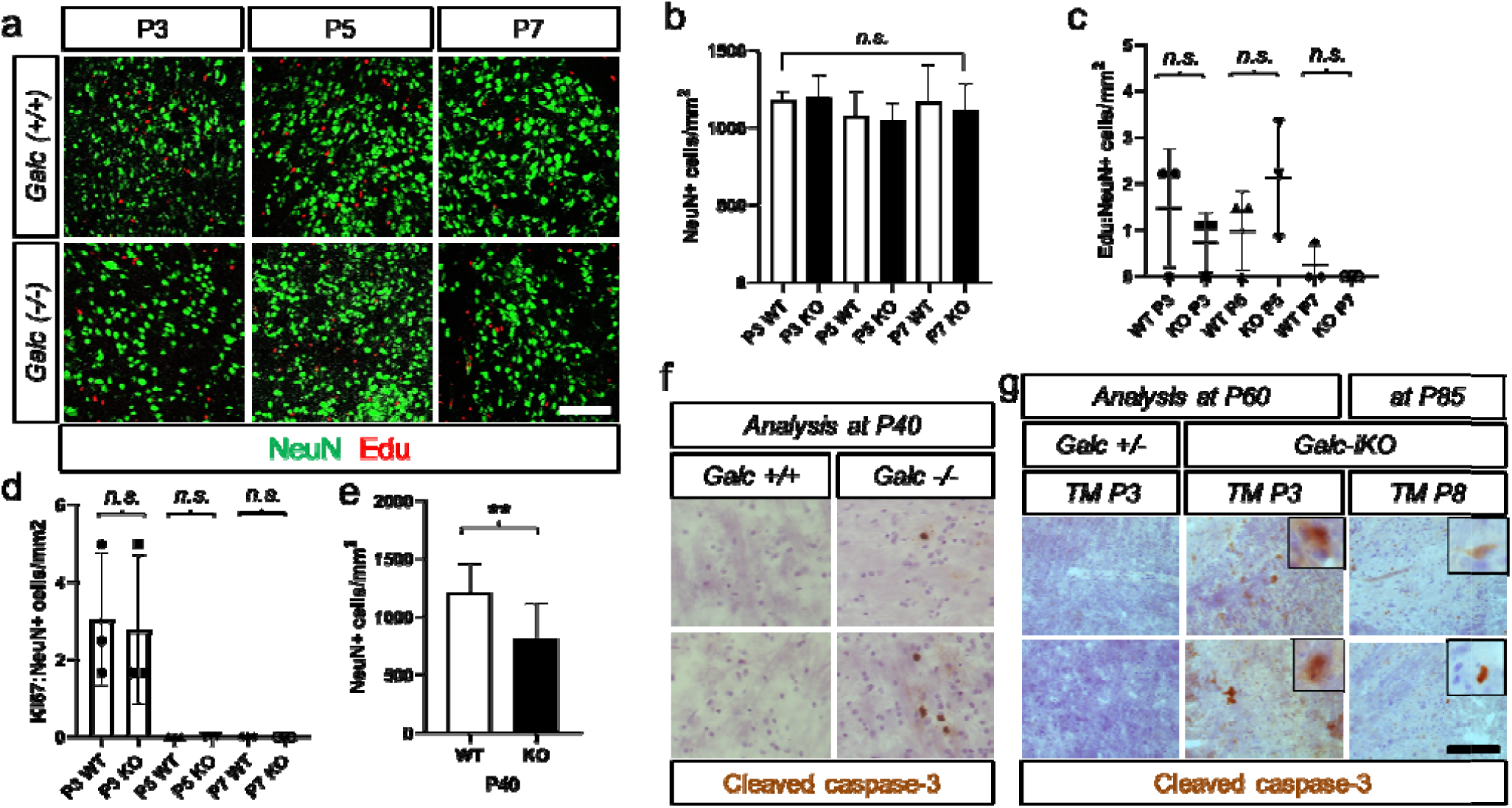
The moribund GALC-depleted brainstem had reduced NeuN+ neuron numbers. Immunostaining (**a**) and counting of NeuN+ neurons (**b**) shows that there were not changes in P3, 5, and 7 brainstems of *Galc*-KO. (**c, d**) Proliferating NeuN+ cell numbers were also not changed during this period. Edu was administrated 24 hours before of the analysis. (**e**) The end-stage KO brainstem has a reduction of NeuN+ cell number significantly. Scale bar=50 *µ*m. Immunostaining of cleaved caspase-3 revealed that apoptotic cell death is significantly increased in the brainstem of moribund *Galc-*KO (**f**) and *Galc*-iKOs (**g**). *Galc*-iKO≤P4 had more prominent signals than *Galc*-iKO≥P6. Scale bar=100 *µ*m.

**Figure S4.**
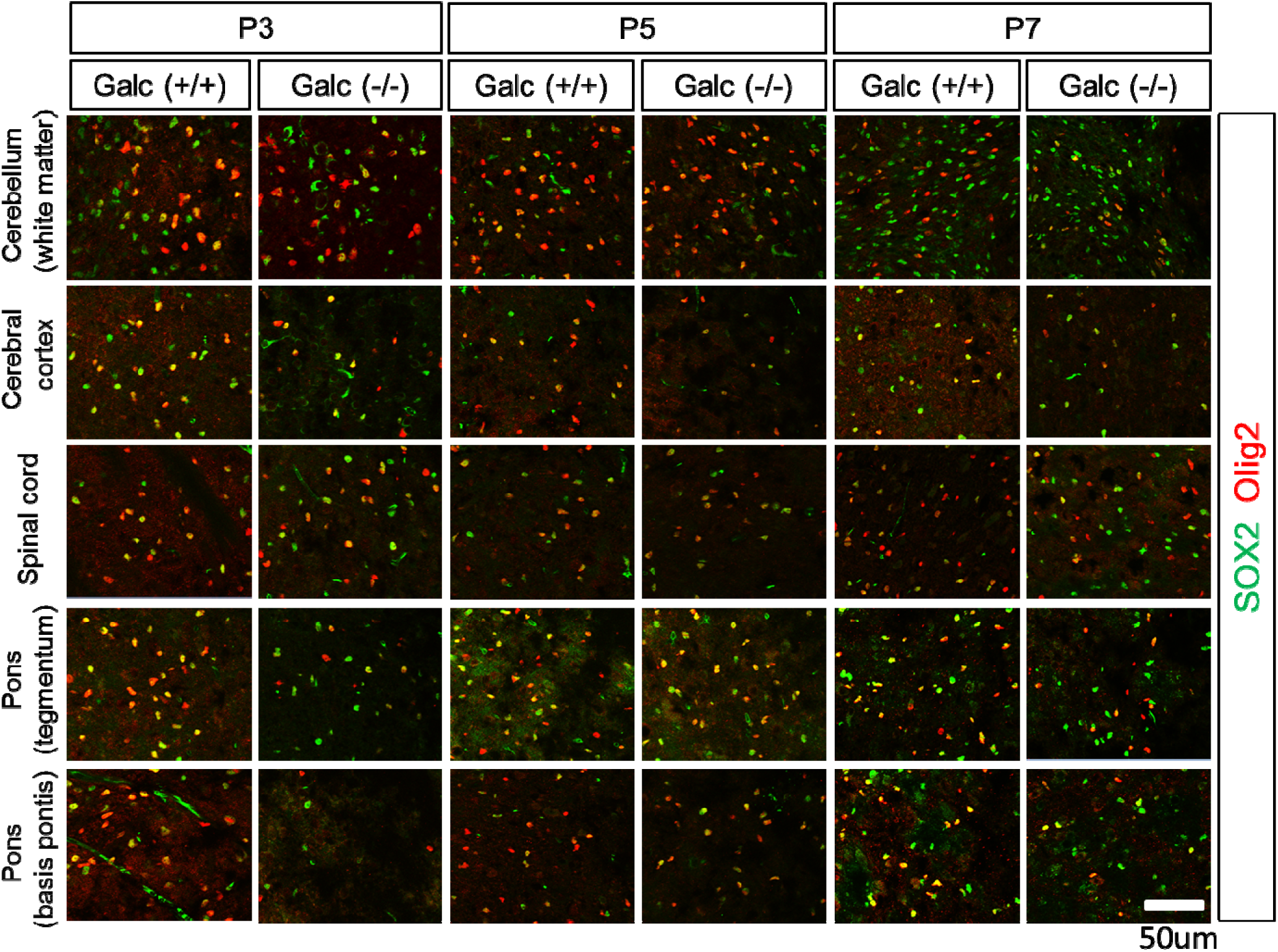
Representative images of SOX2 (green) and Olig2 (red) staining in the brains of *Galc*-WT and KO at P3, P5 and P7.

**Figure S5.**
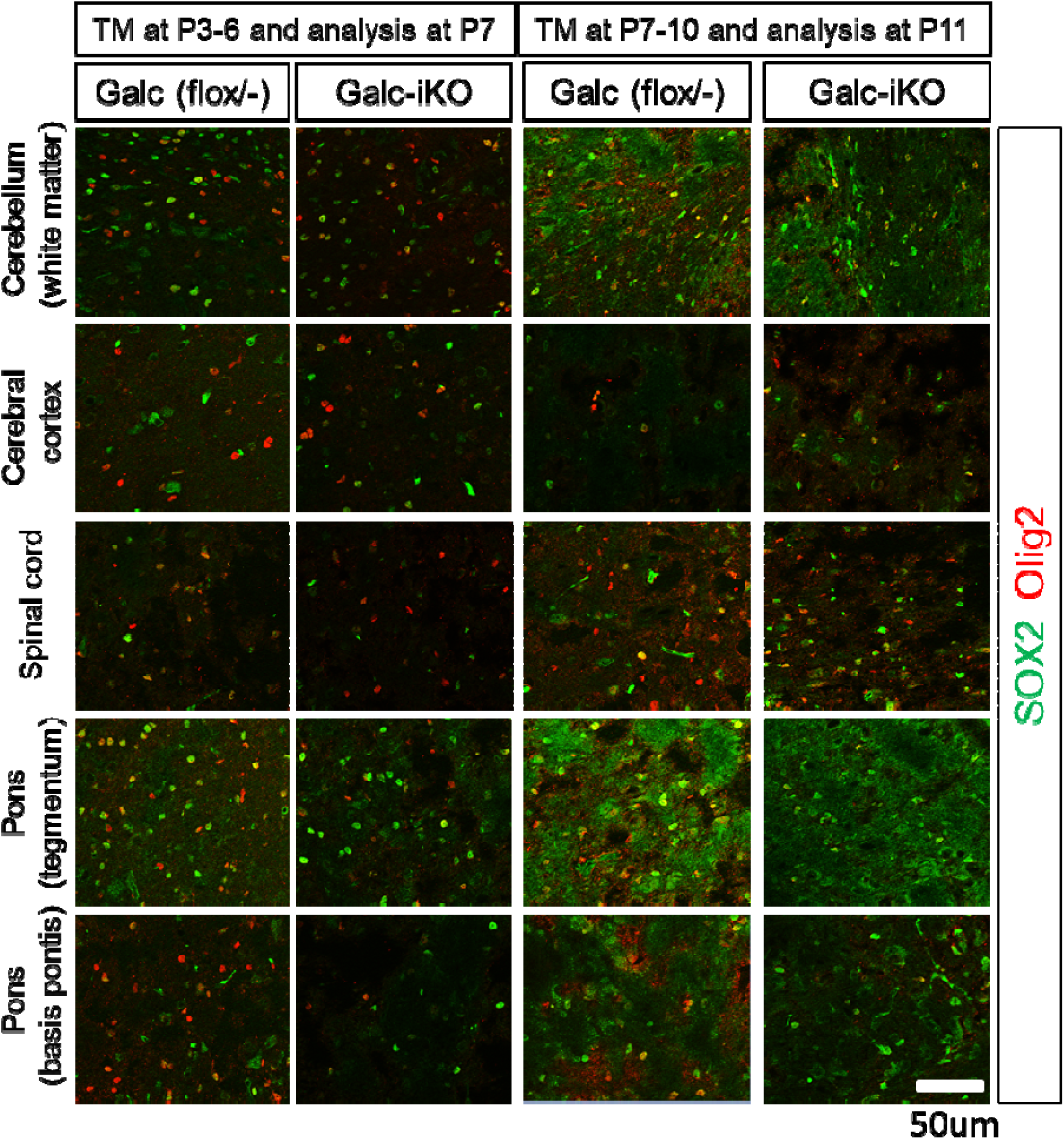
Representative images of SOX2 (green) and Olig2 (red) staining in the brains of *Galc* iKO mice 24 hours after induction starting at either P3 or P7.

**Figure S6.**
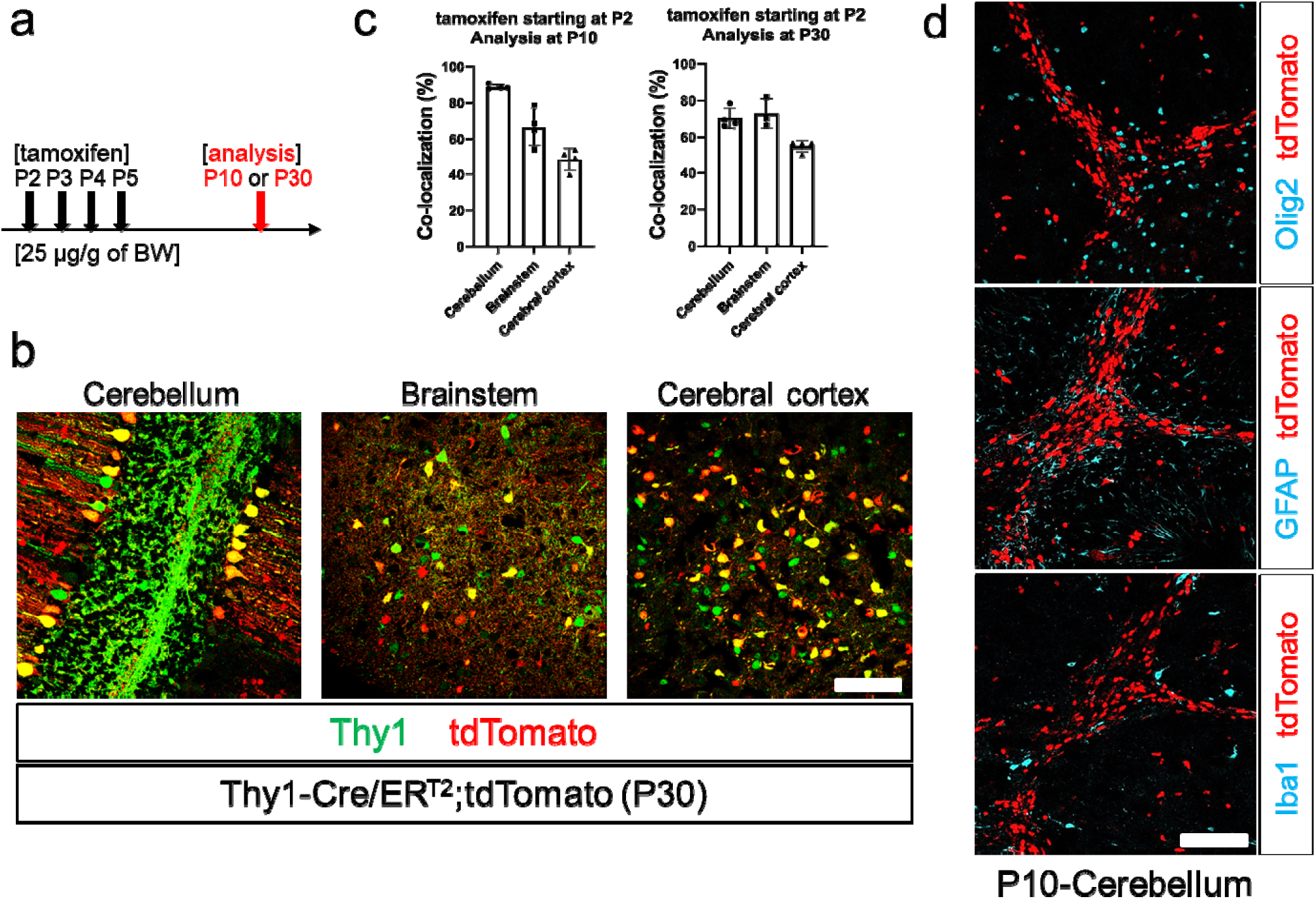
Thy1-Cre/ER^T2^ transgenes specifically undergo recombination in neurons. Tamoxifen was injected into Thy1-Cre/ER^T2^; tdTomato starting at P2, and the mice were dissected at P10 or P30 (**a**). Analysis of major sub-brain regions such as cerebellum, brainstem and cerebral cortex (**b**) showed that more than 65% of Thy1-positive cells overlap with the tdTomato in the hindbrain. The co-localization rate was about 50% in the cerebral cortex (**c**). The tdTomato signals were not overlapped with other cell markers, Olig2, GFAP and Iba1, indicating neuron-specific recombination (**D**). Scale bar=50 *µ*m.

